# Adaptive tail-length evolution in deer mice is associated with differential *Hoxd13* expression in early development

**DOI:** 10.1101/2021.12.18.473263

**Authors:** Evan P. Kingsley, Emily R. Hager, Jean-Marc Lassance, Kyle M. Turner, Olivia S. Harringmeyer, Christopher Kirby, Beverly I. Neugeboren, Hopi E. Hoekstra

## Abstract

Variation in the size and number of axial segments underlies much of the diversity in animal body plans. Here, we investigate the evolutionary, genetic, and developmental mechanisms driving tail-length differences between forest and prairie ecotypes of deer mice (*Peromyscus maniculatus*). We first show that long-tailed forest mice perform better in an arboreal locomotion assay, consistent with tails being important for balance during climbing. The long tails of these forest mice consist of both longer and more caudal vertebrae than prairie mice. Using quantitative genetics, we identify six genomic regions that contribute to differences in total tail length, three of which associate with vertebra length and the other three with vertebra number. For all six loci, the forest allele increases tail length, consistent with the cumulative effect of natural selection. Two of the genomic regions associated with variation in vertebra number contain Hox gene clusters. Of those, we find an allele-specific decrease in *Hoxd13* expression in the embryonic tail bud of long-tailed forest mice, consistent with its role in axial elongation. Additionally, we find that forest embryos have more presomitic mesoderm than prairie embryos, and that this correlates with an increase in the number of neuromesodermal progenitors (NMPs), which are modulated by Hox13 paralogs. Together, these results suggest a role for *Hoxd13* in the development of natural variation in adaptive morphology on a microevolutionary timescale.

**HIGHLIGHTS:**

- In deer mice, the long-tailed forest ecotype outperforms the short-tailed prairie ecotype in climbing, consistent with the tail’s role in balance.
- Long tails are due to mutations on distinct chromosomes that affect either length or number of caudal vertebrae.
- QTL mapping identifies Hox clusters, one gene of which – *Hoxd13 –* shows low allele-specific expression in the embryonic tail bud of forest mice.
- Forest mouse embryos have a larger presomitic mesoderm (PSM), likely mediated by a larger progenitor population (NMPs) and lower *Hoxd13* levels.

## INTRODUCTION

Understanding the genetic and developmental bases of evolutionary changes in morphology, especially those that affect fitness in the wild, is a key goal of modern biology (Carroll 2000, 2008, Arthur 2002, Gilbert & Epel 2009). A major source of morphological change on a macroevolutionary scale in animals is the alteration in the numbers and identities of serially homologous body parts along the anterior-posterior axis – from body segments of arthropods and annelids to vertebrae in the spinal column of vertebrates. Much work has been done to understand the mechanistic basis of changes in segment identity, for example, how shifts in the expression profiles of developmental genes are associated with large-scale changes in the body plan of invertebrates (Averof & Patel 1997, Liubicich et al. 2009) and with transposition of vertebral identities in vertebrates (Burke et al. 1995). However, relatively little is known about how genetic changes act through developmental processes to produce differences in segment size and/or number that occur in nature, and if the same mechanisms involved in macroevolutionary changes contribute to variation within or between closely related species.

In vertebrates, segment identity and number are determined embryonically. During the process of main body axis segmentation, the embryonic segments – somites – form rhythmically from anterior to posterior as the embryo elongates. As somite formation proceeds, the unsegmented presomitic mesoderm shrinks, and segmentation ends when somite formation catches up to the tip of the elongating tail (Bellairs 1986, Gomez & Pourquié 2009, Mallo 2020). Periodic expression of Notch pathway component regulates the rate of segment formation (Gomez et al. 2008, Schröter & Oates 2010, Harima et al. 2013), and posterior axis elongation is promoted by Wnt and FGF activity in the tail bud (Aulehla & Pourquié 2010). Changes to the dynamics of somite formation and/or posterior elongation are thought to largely underlie evolutionary differences in segment number (Gomez & Pourquie 2009). Concomitantly, regionalized morphologies of axial segments are influenced by expression domains of Hox genes, the boundaries of which correlate to regional vertebral identity (Kessel & Gruss 1991, Burke et al. 1995, Wellik 2007, Mallo et al. 2010). The role of Hox genes in conferring segmental identity are complemented by their role in regulating axial growth. In particular, activation of posterior Hox genes correlates with a slowdown of axis elongation via the repression of Wnt activity (Young et al. 2009, Denans et al. 2015, Diaz-Cuadros et al. 2021).

In vertebrates, one of the most variable segmental morphologies is vertebra number, especially those in the tail. In mammals, the number of cervical vertebrae is nearly uniform: the vast majority of mammals have seven cervical vertebrae with a few well-known exceptions (Asher et al. 2011, Varela-Lasheras et al. 2011, Buchholtz 2012). In contrast, the caudal region is the most evolutionarily labile region of the vertebral column, ranging from as few as three vertebrae in the coccyx of great apes to more than 45 in the long-tailed pangolin (Flower & Lydekker 1891, Buchholtz 2012). Tail morphology is often closely associated with its function – from propulsion during swimming (Fish 2016), a counterweight during bipedal saltation (O’Connor et al. 2014) or as a rudder during gliding (Essner 2002) or powered flight (Lawlor 1973) – suggesting a role for natural selection in the evolution of the tail.

The deer mouse (*Peromyscus maniculatus*) occupies diverse habitats across its extensive range in North America and shows striking variation in several morphological traits, most notably, tail length (Osgood 1909, Dice 1940, Blair 1950). At the extreme, deer mice occupying forest habitat can have tails that are 60% longer (approximately 45 mm difference) than those occupying prairie habitat (Kingsley et al. 2017). Remarkably, this morphological divergence between the forest and prairie ecotypes evolved recently, likely as a result of the northward retreat of glaciers approximately 10,000 years ago that opened up new forest habitats, which prairie mice could colonize and where selection may have favored the evolution of long tails (Osgood 1909, Kingsley et al. 2017). Indeed, in this species, long tails may be beneficial for arboreal locomotion: long tails have evolved multiple times independently in forested habitat (Kingsley et al. 2017); tail amputation adversely affects climbing performance, disproportionately reducing performance in forest mice (Horner 1954); and specifically, longer tails are predicted to more effectively promote balance than short tails (Hager & Hoekstra 2021).

Here, we investigate the potential behavioral consequences and the genetic and developmental causes for natural differences in tail length by comparing two representatives of classic deer mouse ecotypes – *P. m. nubiterrae* (forest) and *P. m. bairdii* (prairie) (Osgood 1909) – found in eastern North America (Fig. 1A, B). First, we show that these two subspecies differ dramatically in their climbing performance, in the direction expected based on their tail length differences. Then, using a forward-genetics approach, we identify regions of the genome harboring mutations that affect tail length. We link changes in expression of a gene in one of these regions, *Hoxd13*, to differences in presomitic mesoderm size and its neuromesodermal progenitors as a likely developmental mechanism underlying vertebra number differences. Together, these data suggest a role for Hox genes in microevolutionary changes underlying natural variation in morphology.

**Figure 1.**
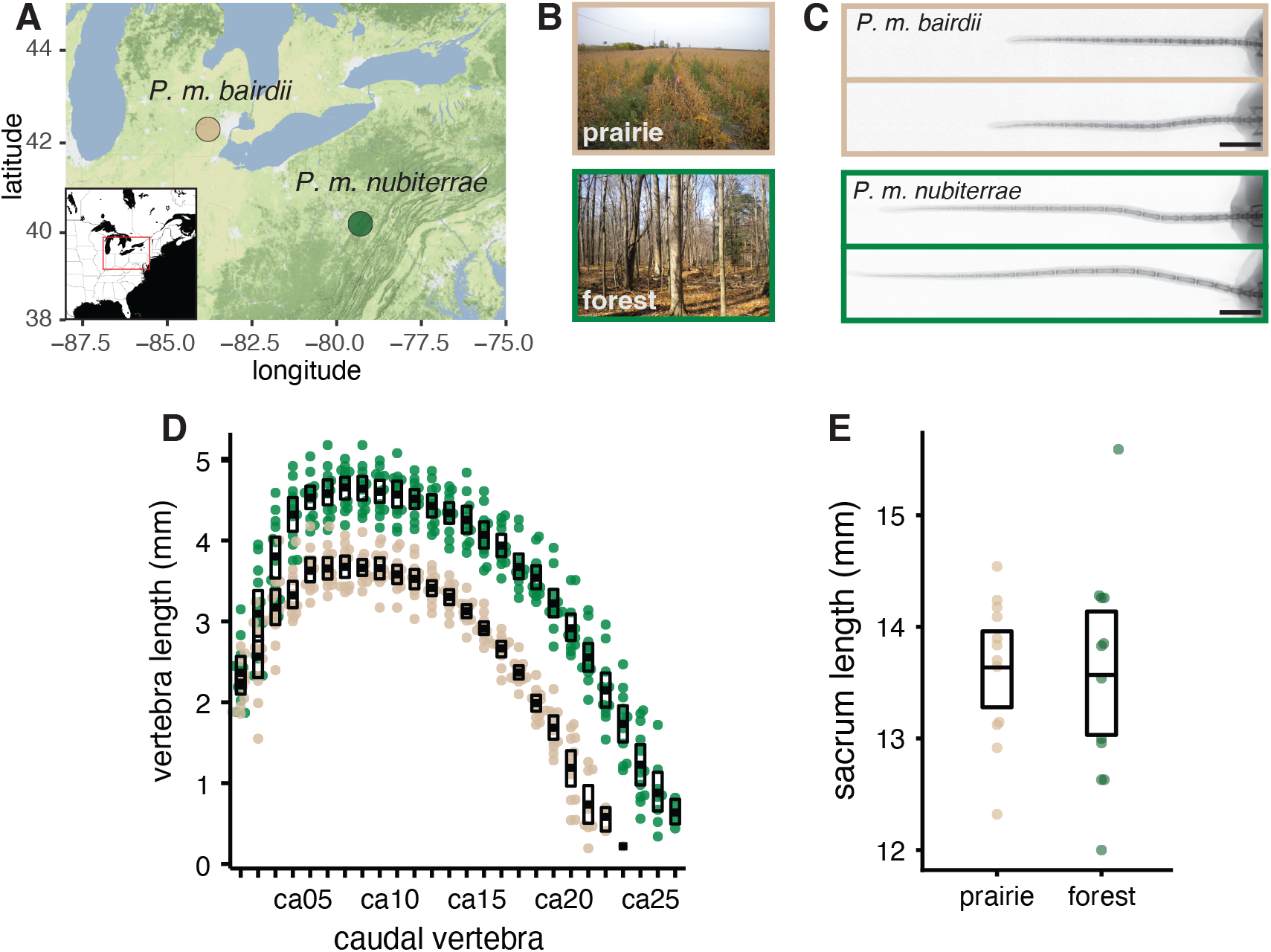
Source populations and morphological traits of wild-caught, laboratory-reared deer mice. Prairie ecotype (*P. m. bairdii*, tan), forest ecotype (*P. m. nubiterrae*, green). **A**. Terrain map showing the trapping locations of mice used in this study: southern Michigan (prairie) and northwestern Pennsylvania (forest). **B**. Photographs represent typical habitat of each ecotype. **C**. Representative radiographs of lab-born prairie (top, n = 2) and forest (bottom, n = 2) mouse tails show differences in tail length. Scale bar = 10 mm. **D**. Scatter plot of caudal vertebra lengths shows that both length and number of caudal vertebrae contribute to differences in tail length between prairie (n = 12, tan) and forest (n = 12, green) mice. **E**. Plots show sacral length, a proxy for body size, does not differ between ecotypes.

## RESULTS

### Tail-length difference due to variation in both caudal vertebral length and number

To characterize the difference in tail length between ecotypes, we measured total tail length, caudal vertebra lengths, and caudal vertebra number from x-ray images of lab-raised forest and prairie mice (n = 12 for each ecotype; Fig. 1C, Fig. S1). We found that forest mice have tails that are 1.4 times longer than those of prairie mice (forest, mean tail length: 84.5 mm [standard deviation (SD): 7.07]; prairie: 60.2 mm [SD: 3.51]), which largely recapitulates the difference observed in wild-caught specimens (1.5-fold difference; Kingsley et al. 2017). Because this difference was maintained when mice were raised in a common environment, variation in tail length likely has a strong genetic (i.e., inherited) component. Specifically, we estimated that genetic variants segregating between ecotypes could explain as much as 88% of the total variance in tail length, based on the distribution of tail lengths in mice from our laboratory colonies.

The difference in overall tail length was due to a difference in both length of caudal vertebrae and number of vertebrae. Because the lengths of vertebrae along the tail of an individual were highly correlated (mean correlation between neighboring vertebrae = 0.86; Fig. S2), hereafter, we focus on the length of the longest vertebra. We found that forest mice have longer caudal vertebrae than prairie mice: the mean length of the longest forest vertebra was significantly longer than that in prairie mice (1.23 times longer; t-test, t = -4.3, df = 7.4, *p* = 2e-3; forest, 4.73 mm [SD: 0.27]; prairie, 3.75 mm [SD: 0.23]). In fact, nearly half of the vertebrae in the forest tail (12 positions, ca4–ca16) are longer, on average, than any vertebra in the prairie tail (Fig. 1D). By contrast, we did not find length differences between ecotypes in vertebrae from other, more cranial regions (e.g., sacral vertebrae; Fig. 1E, Fig. S1). In addition, forest mice have, on average, approximately four additional caudal vertebrae (mean vertebra number: forest, 25.1 [SD: 0.8]; prairie, 21.2 [SD: 0.9]), but no difference in vertebra number in other body regions (Kingsley et al. 2017). Together, a model including only variation in longest vertebra length and vertebra number accounts for nearly all of the variation in total tail length (*R*^*2*^ = 0.99, P < 0.001; *total ∼ longest + number*). Moreover, vertebral length and number contribute approximately equally to the overall tail-length difference between forest and prairie mice (Fig. 1D; Kingsley et al. 2017). Together, these data show that heritable differences in total tail length between forest and prairie ecotypes are due to differences in both the length and number of the constituent caudal vertebrae.

### Forest mice perform better than prairie mice in an assay of arboreal locomotion

The repeated association between long tails and forest habitat suggests an adaptive role for the mammalian tail in arboreal lifestyles in mammals, generally (e.g., Mincer & Russo 2020) and deer mice, specifically (Osgood 1909, Dice 1940, Blair 1950, Kingsley et al. 2017). Indeed, recent models suggest that a longer tail relative to body size is relevant for balance (i.e., controlling body roll) during arboreal locomotion in diverse species (Jusufi et al. 2010, Fukushima et al. 2021, Hager & Hoekstra 2021). Thus, theory predicts that long-tailed forest mice will perform better than similarly-sized prairie mice in behaviors typical of an arboreal lifestyle. To test this prediction in these subspecies, we used a horizontal rod-crossing assay designed to mimic small-branch locomotion (Fig. 2A). We tested the performance of naive adult mice (forest, n = 32; prairie, n = 31) by measuring how often the mice fell from the narrow (0.4 cm diameter) rod and whether they crossed the full length of the rod (44 cm) to another platform (“completed” a cross) (Fig. 2B, Video S1, S2). Forest mice were much less likely to fall: the probability of a forest mouse falling on a given cross is 0.7% (logistic mixed effects model: odds = 0.0073:1), nearly 70 times less than a prairie mouse (48%; odds = 0.906:1; *p* = 7e-9) (Fig. 2C). On attempts when a mouse did not fall, forest mice were much more likely to complete a cross (e.g., ten times more likely on the first cross; logistic mixed effects model: baseline forest probability of completion 72%, odds = 2.5:1; prairie probability 7.3%, odds = 0.08:1; *p* = 8e-8) (Fig. S3). Thus, we find that long-tailed forest mice, even after being reared in laboratory conditions and without prior climbing experience, perform better in this rod-crossing assay than short-tailed prairie mice, consistent with a role for tail-length differences in arboreal adaptation in these subspecies.

**Figure 2.**
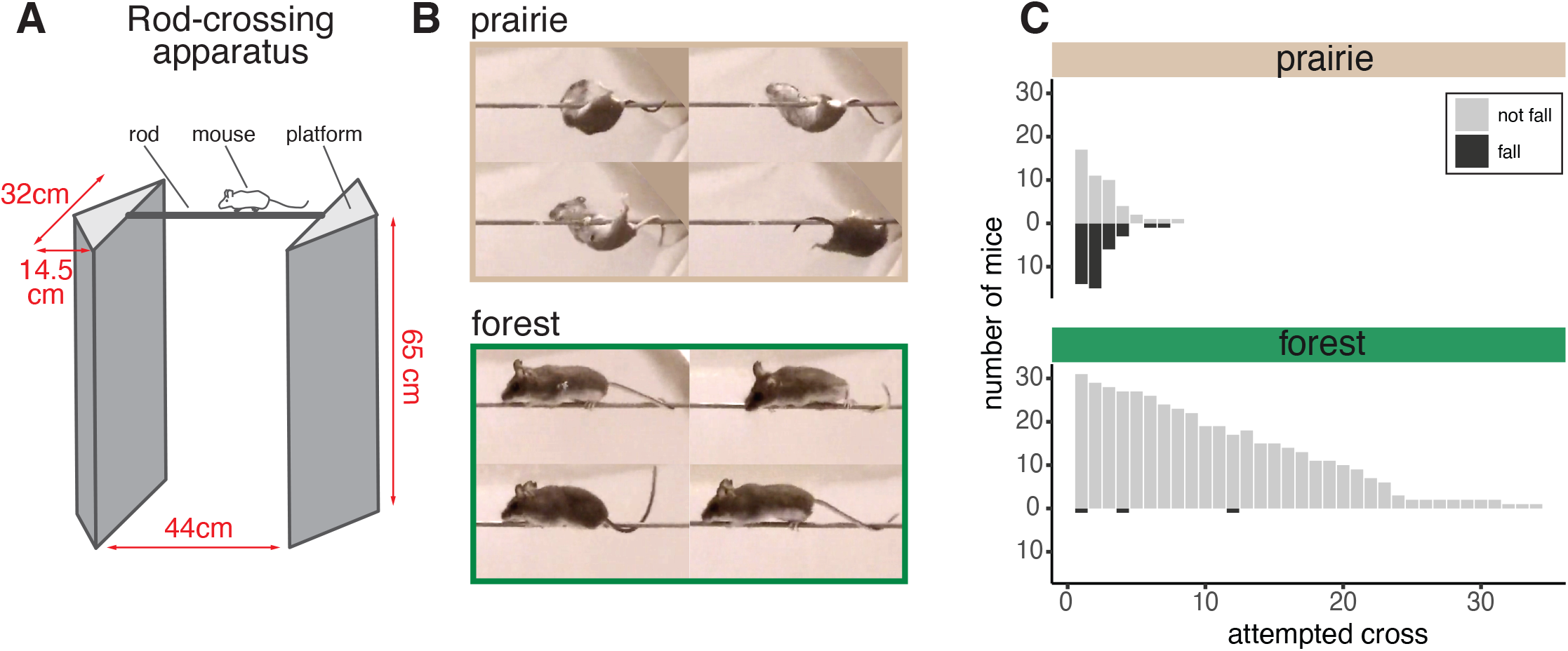
Difference in climbing performance between prairie and forest ecotypes. **A**. Schematic of the rod-crossing apparatus including dimensions (red). **B**. Representative side-view images captured from videos of a rod-crossing assay: prairie (top) and forest (bottom) mice. See Video S1, S2. **C**. Number of mice that fell (dark gray) or did not fall (light gray) on each attempted cross. Prairie mice (top, n = 31) fall more often (p = 1e-12) than forest mice (bottom, n = 32).

### Genetic mapping localizes genomic regions contributing to tail length variation

To characterize the genetic architecture of tail-length variation, we generated a reciprocal genetic cross between forest and prairie mice (n = 4 parents; 1 male and 1 female of each ecotype), resulting in 28 F1 hybrids, which then were intercrossed to produce 495 second-generation (F2) hybrids. Based on the ecotypic differences and trait correlations in the hybrids, we focused on three tail traits for genetic dissection: total tail length, length of the longest caudal vertebra, and number of caudal vertebrae (Fig. 3A, B). The two length traits correlated strongly with body size (Fig. S2), so we used sacrum length as a proxy for body size to adjust values in all subsequent analyses of these traits (see Methods). In the F2 hybrid mice, vertebra length and vertebra number are both significantly correlated with total tail length (Fig. 3C, D, Fig. S2): a linear model with vertebral length and number as the explanatory variables accounts for almost 90% of the variance in total tail length in the F2 hybrid population (*R*^*2*^ = 0.89). However, vertebral length and number were only weakly correlated with each other (*r* = 0.16, p < 0.01; Fig. 3D), suggesting that variation for these two traits is genetically separable. For all three focal traits, F1 hybrid trait values were intermediate between the means of the parental traits (Fig. S1), and F2 trait values fell within the mean parental trait values (Fig. 3B). However, for all three tail traits, a few F2 hybrids had trait values similar to the parental phenotypes, consistent with the trait variation being largely oligogenic (Fig. 3B), making these traits amenable to genetic dissection.

**Figure 3.**
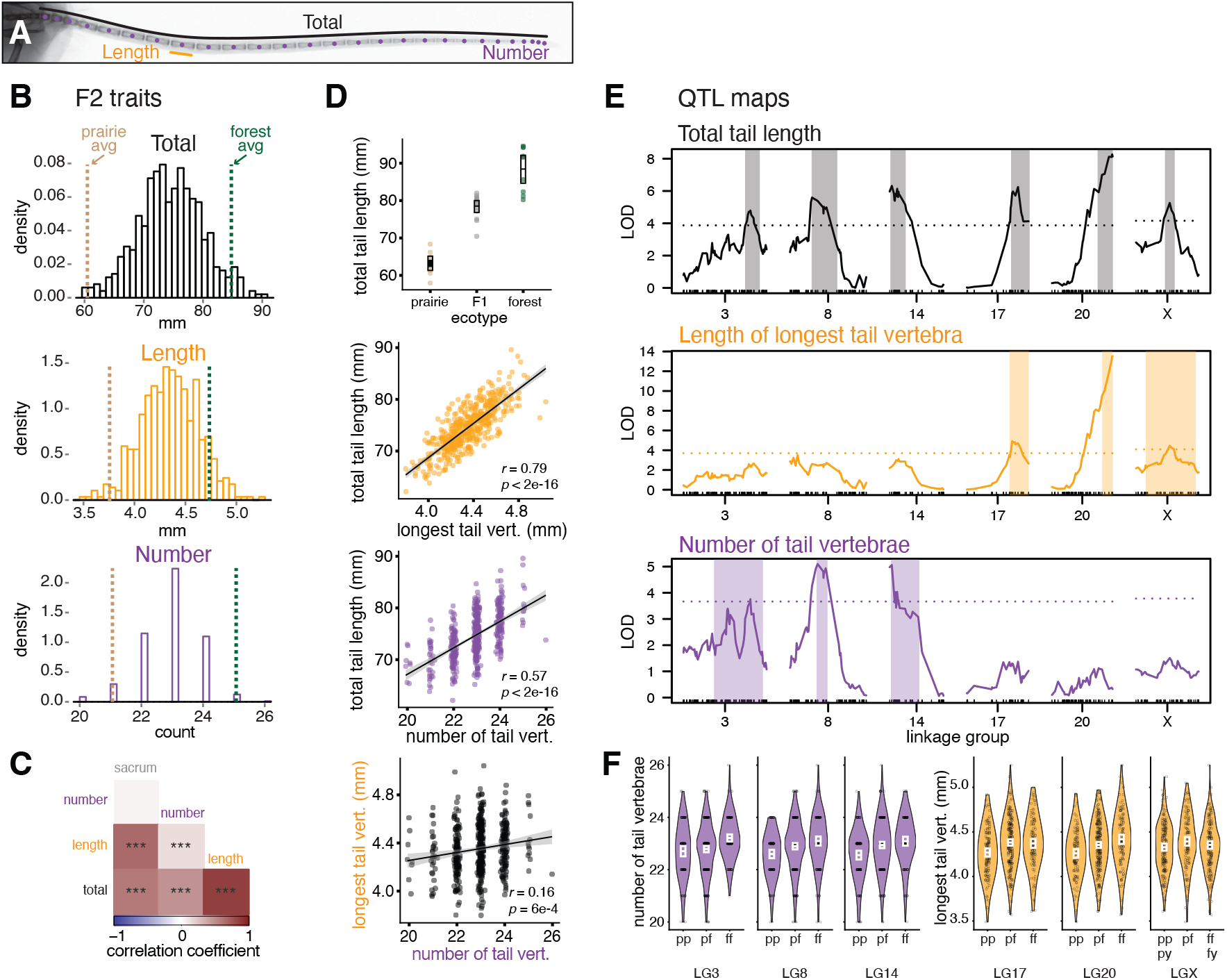
Quantitative trait locus (QTL) mapping in a forest-prairie F2 intercross for three tail traits. **A**. Tail x-ray highlighting focal measurements: total tail length (black), length of the longest vertebra (orange), and number of vertebrae (purple). **B**. Distributions of tail traits in F2 hybrid mice (n = 495). Dashed vertical lines indicate parental trait means: forest (green) and prairie (tan). **C**. Pairwise Pearson correlations among tail traits and sacrum (a proxy for body size). *** indicates p < 0.001. **D**. Plot showing total tail length in each ecotype and their F1 hybrids. Boxes show mean and bootstrapped 95% confidence limits of the mean (top). Scatter plots showing the pairwise relationship between the three tail traits in F2 hybrid mice (bottom 3 plots). **E**. Statistical association (LOD, or log of the odds, score) showing significant QTL associations on six linkage groups for total tail length (top, black); length of the longest caudal vertebra (middle, orange); the number of caudal vertebrae (bottom, purple). Shaded rectangles delineate the Bayesian credible interval (0.95 probability coverage) for each significant QTL. Dotted lines indicate genome-wide significance thresholds (p = 0.05) as determined by permutation tests. **F**. QTL effects on vertebra number (purple, left 3 plots) and vertebra length (orange, right 3 plots) by genotype (pp = homozygous for the prairie allele, pf = heterozygous, ff = homozygous for the forest allele; py = hemizygous prairie male, fy = hemizygous forest male) at the peak LOD marker for each QTL. White boxes show means and bootstrapped 95% confidence limits of the mean.

We next used interval mapping to localize regions of the genome that influence variation in tail traits in our F2 hybrid population. For total tail length, we identified six significant quantitative trait loci (QTL) that, together in a multiple-QTL model, explain 23.8% of the variance in tail length (Fig. 3E, Table S1). The 95% Bayesian confidence intervals (CI) for three of these QTL coincided with those for the three QTL associated with the length of the longest caudal vertebra, which together explained 14.0% of the variance in vertebra length (Table S1). The remaining three QTL for total tail length coincided with three QTL that influence the number of caudal vertebrae; these QTL explained 11.7% of the variance in vertebra number (Table S1). We also identified two additional weak associations for vertebra length (linkage groups 13 and 21), but they did not overlap with QTL for total tail length or vertebra number (Fig. S4, Table S1). The distribution of QTL for vertebral length and number on separate chromosomes conforms with the weak correlation between these traits, consistent with vertebral length and number being under independent genetic control.

By examining the effects of each QTL, we estimated the dominance patterns of each allele. We found that alleles inherited from the forest parent exhibit incomplete dominance (Fig. 3F, Table S1), with varying degrees of mean dominance-effect estimates ranging from -0.06 to 1.55 (where -1, 0, and 1 correspond to complete recessivity, additivity, and complete dominance, respectively; *d/a*, Falconer & Mackay 1996). In a multiple-QTL model, additive effects of forest alleles at the three vertebra-length QTL ranged from 0.02 mm to 0.10 mm, while the additive effects of forest alleles at vertebra number QTL were nearly equal (0.26 to 0.29). Thus, an individual with all three forest alleles at the vertebra length QTL had, on average, a 0.32 mm longer vertebra, and at the three vertebra-number QTL had an average of 1.66 more vertebrae, than an animal with prairie alleles at the respective loci. Together these major-effect QTL explained approximately 33% of the difference of mean vertebra length and 43% of the mean vertebra number difference between forest and prairie ecotypes.

Finally, we performed a sign test that assesses if the direction of the allelic effects at multiple QTL differ from random expectations (Orr 1998, Fraser 2020). We found that for each of the six QTL associated with total tail length, the forest allele effect was always in the expected direction (Fig. 3F), that is, forest alleles result in larger trait values, a pattern that deviates from neutral expectations (*p* = 0.045; Orr’s QTLSTEE, see Methods). In addition, a test for directional selection based on the ratio of parental and F2 trait variances also departs significantly from the neutral expectation (*v* = 9.7, *p* < 0.01; *p* < 0.05 for H^2^ < 0.73; Fraser 2020). These observations provide additional, independent support for the hypothesis that natural selection favors longer tails in forest deer mice.

### Variation at the *Hoxd13* locus is associated with differences in caudal vertebra number

The striking divergence in caudal vertebra number we identified between ecotypes provided an opportunity to explore the genetic and developmental mechanisms that lead to intraspecific segment number evolution. We therefore decided to focus on one tail measure – vertebra number – for further investigation. The number of caudal vertebrae is established *in utero* (Fig. S5). Therefore, to aid in the prioritization of potentially causative genes and to better understand the developmental pathways likely to be important in establishing the vertebra number difference between these ecotypes, we first performed RNA-seq on tail bud tissue spanning the period in which tail somites are forming (“early”, E12.5 to “late”, E15.5, which correspond to E10.5 and E13.5 in *Mus musculus*; Theiler 1989, Manceau et al. 2011, Davis & Keisler 2016) to identify genes that are differentially expressed, even at low levels, between ecotypes (forest, n = 18; prairie, n = 17). In a multidimensional scaling analysis, these samples clustered strongly both by ecotype (forest/prairie) and by stage (early/late tail segmentation) (Fig. S6A). By comparing expression levels between ecotypes, we found 2534 and 3467 protein-coding genes in early and late stages, respectively, that were differentially expressed between forest and prairie embryonic tails (FDR-adjusted p < 0.05; Fig. S6B). Of these, 1515 were differentially expressed in the same direction in both stages, while 1017 were differentially expressed only early on and 1950 only later (2 genes were differentially expressed at both time points, but in opposite directions). Thus, perhaps not surprisingly, we found thousands of genes differentially expressed between these ecotypes during a window critical for somitogenesis.

Variants that are causative for the difference in vertebra number are expected to lie within the three relevant QTL confidence intervals. We therefore next identified the annotated protein-coding genes within each QTL confidence region (n = 527, linkage group [LG] 3; n = 85, LG8; n = 110, LG14) and intersected these mapping results with the RNA-seq data to identify genes that both fall within QTL confidence intervals and show differential expression. Of the protein-coding genes in these three intervals, we found between 28 and 112 genes in each QTL were differentially expressed during tail development (n = 112, LG3; n = 28, LG8; n = 28, LG14) (Table S2). To identify which of these genes have known effects on tail length, we further prioritized genes that have orthologs with known effects on tail length when manipulated in *Mus* and cataloged in the Mouse Genome Informatics (MGI) Phenotype database (see Methods; Table S3). Of the 155 orthologs included in MGI categories that affect tail length, only five fell within our QTL intervals for vertebra number and also had significant differences in expression levels during embryonic tail elongation: *Sp5, Hoxd13, Hoxd9* (LG3), *Hoxa10* (LG8), and *Apc* (LG14). Hox genes have known roles in axial patterning, and *Sp5* and *Apc* regulate Wnt signaling; thus these genes comprise a list of top candidate genes (Fig. 4A).

**Figure 4.**
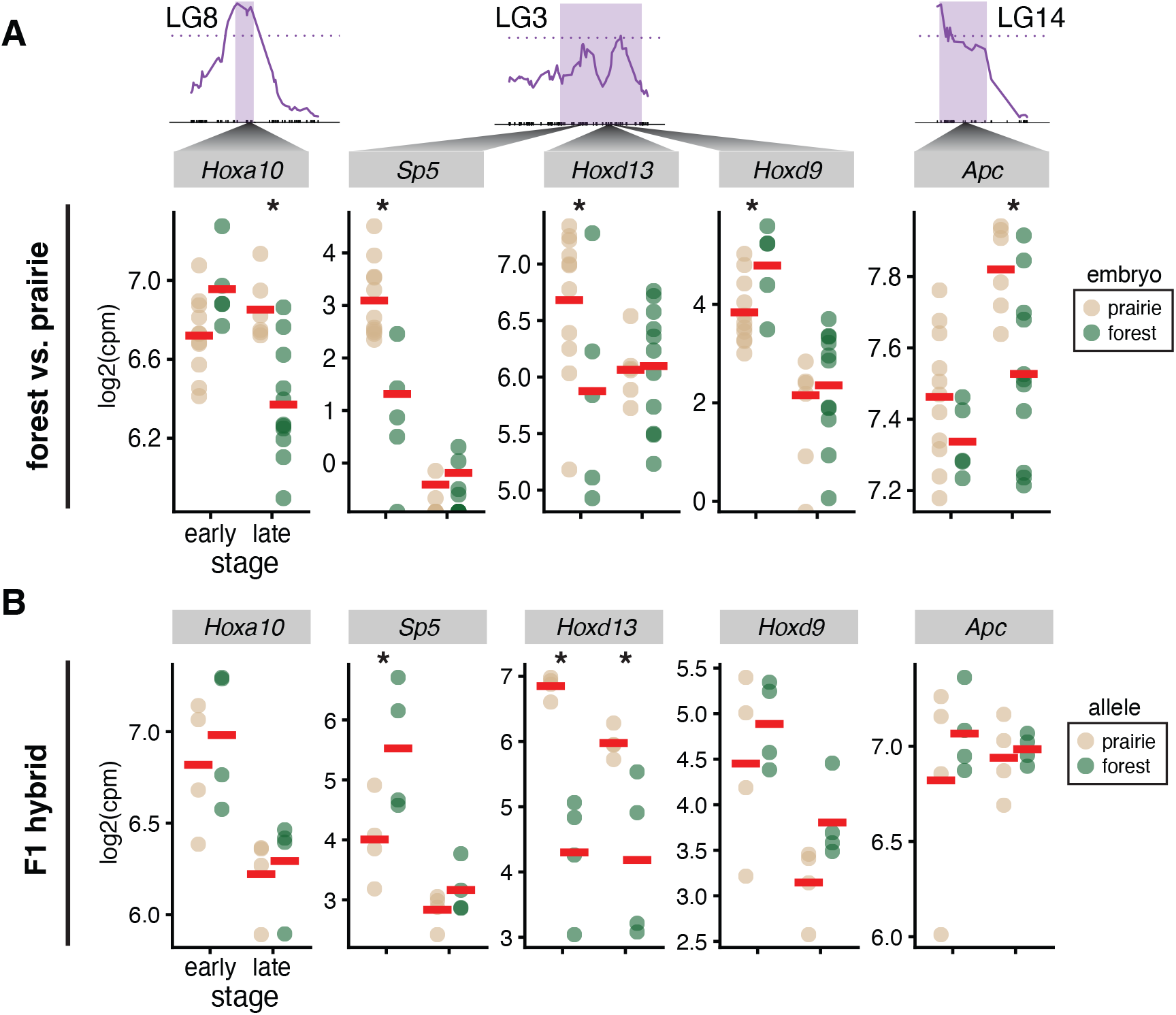
Expression level of five candidate genes associated with vertebra number at two developmental timepoints in embryonic tail from forest and prairie mice. **A**. Top panel: Three vertebra-number QTL with 95% confidence intervals highlighted (purple) that each contain at least one candidate gene. Bottom panel: RNA-seq-estimated gene expression level (cpm = counts per million) for five top candidate genes in forest (n = 18, green) and prairie (n = 17, tan) embryos at early (E12.5–13.5) and late (E14.5–15.5) developmental timepoints. **B**. Allele-specific RNA-seq in F1 forest-prairie hybrid embryonic tails (n = 8). * indicates p < 0.05.

The causal mutations found within QTL regions that affect expression of candidate genes are expected to act in an allele-specific manner (i.e., *cis*-acting). Therefore, we estimated allelic bias in expression using bulk RNA-seq data from F1 hybrid tail bud tissue collected at both early (E12.5) and late (E14.5) tail growth stages (Fig. S7). Of the five candidate genes, only *Hoxd13* showed allele-specific expression differences in the same direction observed between the forest and prairie mice (Fig. 4B). Interestingly, the expression difference between the *Hoxd13* alleles in F1s surpassed the difference observed between ecotypes (log2 fold-change = 0.85 between ecotypes; 2.57 between alleles), suggesting additional *trans-*acting effects that act antagonistically to the *cis*-acting difference. *Hoxd13* has also been shown to be expressed in the tail bud in the laboratory mouse, zebrafish, and lizard during axial elongation (Dollé et al. 1991, Di-Poï et al. 2010, Ye & Kimelman 2020, Guillot et al. 2021). Together, these data point to *cis*-acting mutation(s) that affect the expression of *Hoxd13* in the developing tail as a strong candidate for contributing to differences in caudal vertebra number.

In addition to its expression level, we also compared the entire coding region of *Hoxd13* (1017 bp) between ecotypes. Although mammalian Hox gene sequences are highly conserved (Lin et al. 2008), we found that *Hoxd13* had a 3-bp insertion at amino acid position 109 in the disordered N-terminal region of the protein (Basu et al. 2020). The mutation was fixed between our laboratory colonies of forest and prairie mice (Fig. 5A) and resulted in an expansion of a polyalanine tract from four (forest) to five (prairie) residues; expansions of polyalanine tracts in this region of the protein cause hereditary synpolydactyly in humans (Muragaki et al. 1996, Albrecht et al. 2004). This 3-bp insertion (or 5-alanine tract) is absent in other *Peromyscus* species, *Mus musculus*, and all other rodents we surveyed, and thus appears unique to these prairie mice (*P. m. bairdii*; Fig. 5B).

**Figure 5.**
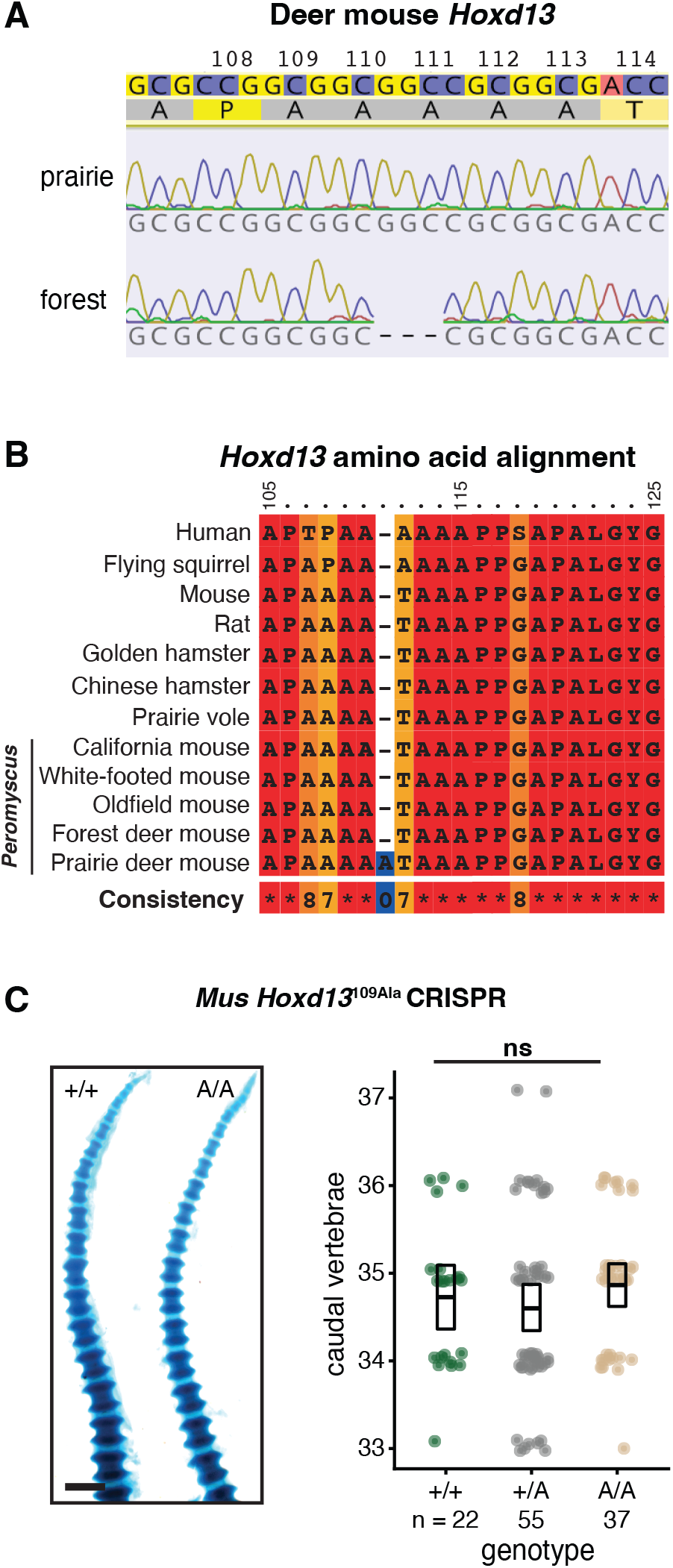
Amino acid variation in Hoxd13 is not associated with vertebra number. **A**. Sequence chromatograms showing a portion of *Hoxd13* exon 1 (positions 108–114aa) aligned to the *P. maniculatus bairdii* reference genome (top). **B**. PRALINE alignment for a portion of the N-terminal region (*Mus* position 105–125aa) of HOXD13 in *Peromyscus*, other rodents and human. **C**. Left panel: Examples of transgenic *Mus* wild type and homozygous for the engineered *Hoxd13* CRISPR allele (109Ala) P0 tails stained with alcian/alizarin. Scale bar = 1 mm. Right panel: Caudal vertebral counts for wildtype (forest genotype, green), heterozygous (gray), and homozygous (prairie genotype, tan) for the 109Ala allele. Boxes show mean and bootstrapped 95% confidence limits of the mean.

We next explored whether this amino acid insertion in *Hoxd13* causes a difference in caudal vertebra number. We first performed a protein variation effect analysis, which predicted that the insertion has a neutral effect on the biological function of the HOXD13 protein (PROVEAN score: 0.561). Next, we used CRISPR-Cas9 mutagenesis in C57BL/6 laboratory mice to introduce an extra alanine residue into the native *Mus* 4-alanine tract at position 109 (*Hoxd13*^109Ala^), thereby replicating the prairie allele in *Mus*. Note that the forest allele encodes a protein identical to the native *Mus* HOXD13. When we intercrossed animals heterozygous for the CRISPR edit and counted the number of caudal vertebrae in second-generation pups at birth (P0; n = 114), we found no significant effect of the alanine insertion on vertebra number: mice that were homozygous for the 109Ala insertion had a mean of 34.9 vertebrae compared to the wild type 34.7 (ANOVA, *p* = 0.41, df = 2; power to detect difference of 0.52 vertebrae at 0.05 significance = 0.72) (Fig. 5C). Together, these results suggest that variation in the *Hoxd13* coding region does not affect vertebra number, and instead points to a change in the *cis*-acting regulation of *Hoxd13* expression during a critical time for tail elongation as a likely genetic mechanism.

### Differences in embryonic tail development correlate with segment number variation

To determine the developmental mechanism contributing to differences in caudal vertebra number, and what role *Hoxd13* may play in deer mice, if any, we compared the developing tail tissues and cell populations of forest and prairie embryos during tail segmentation. Embryonically, vertebrae arise from the sclerotome of the somites, epithelial segments that sequentially bud off at a clock-like rate from the anterior of the presomitic mesoderm (PSM; Christ & Wilting 1992, Dequéant & Pourquié 2008). Segmentation ends – and the number of vertebrae is determined – when somitogenesis catches up to the tip of the growing tail bud (Bellairs 1986, Gomez et al. 2008). Thus, an increase in somite number can be produced by accelerating the rate of somite production (or slowing the progression of the wavefront) resulting in smaller somites, assuming the same rate of posterior elongation, or alternatively, if the rate of somite formation is constant, increasing the size of the PSM (implying a higher rate of PSM production from the tail bud) (Gomez & Pourquie 2009).

To test these hypotheses, we measured the length of both the most-recently-formed somite (S1) and the PSM in E11.5–E15.5 embryos, following the formation of the first post-hindlimb somites (Fig. 6A). We found that S1 lengths did not differ through time between forest and prairie embryos (linear regression, t = 1.28, df = 2, *p* = 0.08; Fig. 6B). Notably, the S1 length differences trended in the opposite direction from expected if somites were produced faster in forest embryos. Moreover, these results are consistent with the rate of somite formation measured in cultured tail bud explants from forest and prairie embryos: we found no significant difference in the rate of somitogenesis (Wilcoxon test: W = 31.5; *p* = 0.45; Fig. S8). By contrast, we found that the length of the PSM was significantly different between ecotypes (linear regression, t = 3.05, df = 2, *p* = 0.004; Fig. 6C), suggesting a different rate of posterior elongation. Specifically, the PSM starts at a similar size but then diverges between ecotypes in the expected direction, that is, larger in forest mice than prairie mice (in embryos with < 6 post-hindlimb somites, there is no significant difference in PSM length [Wilcoxon test, W = 11, p = 1]; for bins 6–12, 12–18, and > 18 somites, forest PSM is an average of 129 µm, 189 µm, and 111 µm longer, respectively, than prairie PSM [Fig. S9]). Thus, by comparing forest and prairie embryos, these results show that the larger number of caudal vertebrae in adult forest mice is likely due to an increased elongation rate, resulting in a longer PSM, rather than an increased rate of somitogenesis.

**Figure 6.**
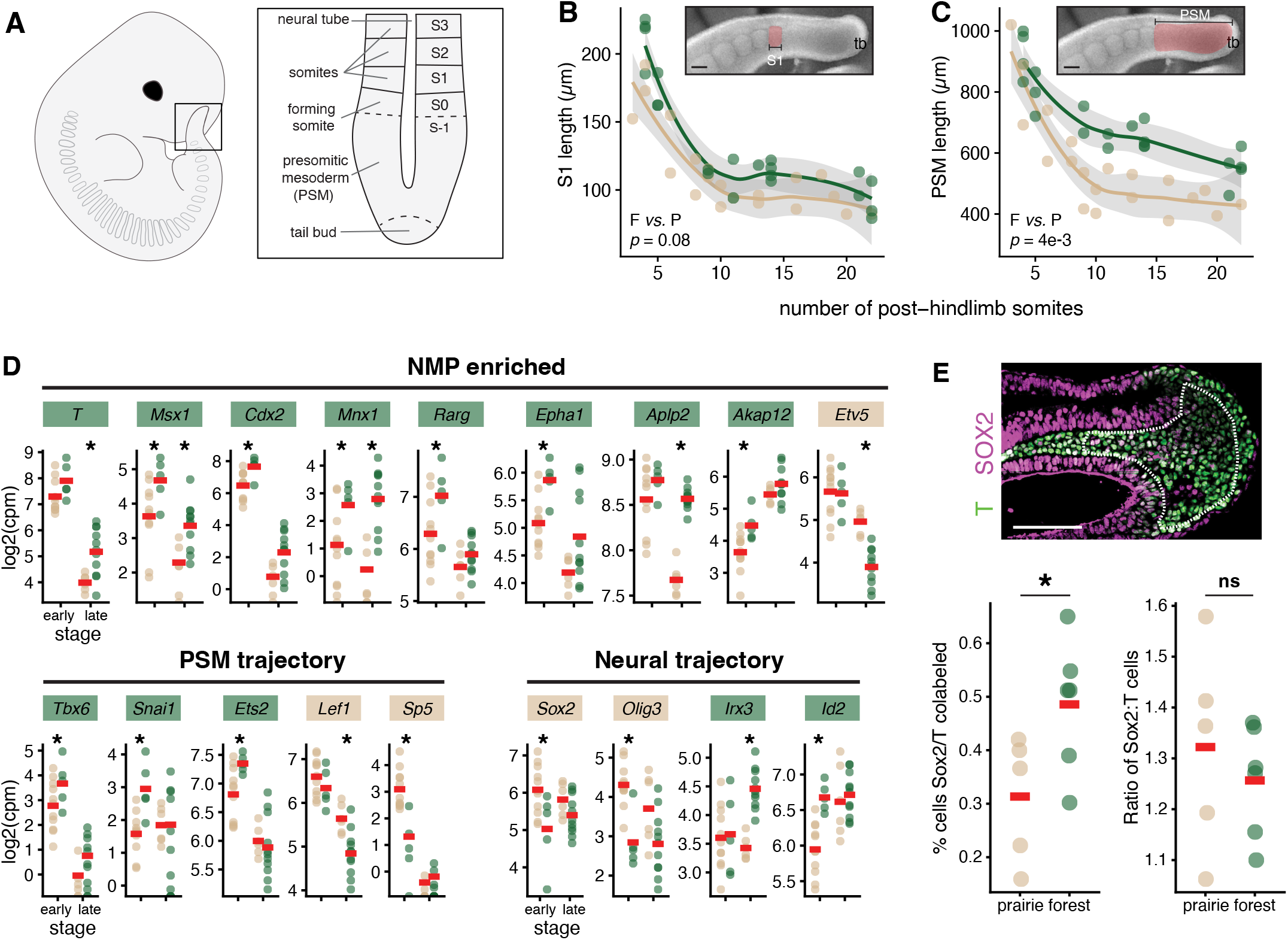
Developmental basis of difference in caudal vertebra number. **A**. Diagram of an E12.5 embryo showing the anatomy of the embryonic tail, including somites (S) and the presomitic mesoderm (PSM). **B**. Length of the most recently formed somite (S1, pink) across tail segmentation stages (E11.5–E15.5, plotted by number of post-hindlimb somites) measured in fixed specimens of forest (n = 20, green) and prairie (n = 18, tan) embryos. tb = tail bud; scale bar = 100 µm. **C**. Length of PSM (pink) measured in fixed specimens across tail segmentation (E11.5–E15.5) in forest (n = 20, green) and prairie (n = 18, tan) embryos. **D**. RNA-seq-estimated transcript counts of genes associated with NMPs as well as PSM and neural fate trajectories that are differentially expressed (adjusted p < 0.05) between forest and prairie embryonic tails at early (E12.5) and late (E14.5) stages of tail development. Gene names are colored according to direction of differential expression (green = higher expression in forest; tan = higher expression in prairie). **E**. Top panel: representative immunofluorescence image from a prairie embryo showing the tail bud mesenchyme (dashed line) in which SOX2 (magenta)- and T (green)-labeled cells were counted. Caudal is to the right. Bottom left: The percentage of co-labelled cells (NMPs) in forest (n = 6, green) and prairie (n = 5, tan) embryonic tail bud sections at E12.5. Bottom right: Ratio of SOX2-labeled cells to T-labeled cells in sections of tail bud mesenchyme. * indicates p < 0.05; Scale bar = 100µm.

Post-anal PSM size is mediated by regulation of a population of bipotential cells in the tail bud that produce the caudal PSM – the neuromesodermal progenitors (NMPs), a cell population in which *Hoxd13* is expressed during tail elongation in *Mus* (Guillot et al 2021). A larger PSM could be produced by either an overall increase in the number of NMP cells or, alternatively, a shift in the balance of NMP fate trajectories towards mesodermal (PSM) to the detriment of neural fates. Indeed, a bias toward the PSM fate in *Mus* results in more segments, whereas a balance tipped toward the neural fate produces fewer (Koch et al. 2017, Aires et al. 2019). To test these alternative hypotheses, we first returned to our transcriptomic data to examine the expression profile of markers enriched in NMP cells as well those for the relevant fate trajectories between forest and prairie mice. Of the genes that were differentially expressed between ecotypes and enriched in NMPs in *Mus* (adjusted *p* < 0.05), eight of nine were more highly expressed in the developing tail buds of forest than prairie mice (Fig. 6D, top), consistent with ecotypic differences in NMP abundance. However, when we examined PSM-versus neural-fate markers, we did not find evidence for a strong shift towards the mesodermal fate. In other words, there was no obvious trend in the genes correlated with the NMP fate trajectories (Fig. 6D, bottom): of the five genes upregulated in PSM trajectory in *Mus* and differentially expressed between ecotypes, three were upregulated in forest mice and two in prairie mice, while of the four genes upregulated in the neural trajectory in *Mus* and differentially expressed, two were upregulated in forest mice and two in prairie mice. Thus, the RNA-seq data suggest that, while there is no clear shift in gene expression associated with two downstream NMP fates (PSM versus neural), the higher expression of NMP-enriched genes is consistent with a larger pool of axial progenitor cells in forest compared to prairie mice.

To confirm a difference in the number of NMP cells between ecotypes, we counted cells in embryonic tail bud sections immunostained for SOX2 and T, canonical markers for NMP cells, at E12.5 (forest, n = 6; prairie, n = 5). We found that a greater proportion of forest tail bud mesenchyme cells are co-labeled with SOX2 and T antibodies than prairie tail buds (t-test; t = -2.4; df = 8.8; *p* = 0.04; Fig. 6E), consistent with the transcriptomic data, indicating a larger pool of axial progenitors in forest ecotype. We also compared the ratio of SOX2:T cells in the tail bud mesenchyme of both ecotypes to test for a bias in NMP fates, with the expectation that long-tailed forest mice would have a lower ratio if NMPs were biased toward producing PSM. However, consistent with the transcriptomic data, we did not detect a significant difference in the ratio of SOX2:T immunostained cells (t = 0.2, df = 7.4, *p* = 0.9; Fig. 6E), although our power to detect a difference was low. Thus, the results from the transcriptomic and immunohistochemistry experiments together suggest that differences in NMP abundance, not a shift in NMP fate dynamics, likely contribute to differences in PSM size between forest and prairie ecotypes.

## DISCUSSION

Here we investigated both the ultimate and proximate mechanisms driving the divergence in a skeletal trait – tail length – between forest and prairie ecotypes within a single species of deer mice. These tail-length differences are due to changes in both caudal vertebral length and number. In the six genomic regions that are associated with tail-length variation, the forest allele is always associated with longer tails, consistent with natural selection driving trait divergence, likely due to longer tails contributing to at least some aspects of climbing performance in forest environments. In one of these genomic regions lies a strong candidate gene, *Hoxd13*, which shows allele-specific differential expression between forest and prairie embryos during tail elongation. These ecotypes also differ in the size of the tissue from which somites develop as well as its underlying progenitor cell population. Taken together, our results suggest a plausible proximate mechanism for the evolution of vertebra number between deer mouse ecotypes: reduced *Hoxd13* expression maintains the progenitor pool of the tail bud PSM in forest mice, leading to prolonged axial extension, the formation of more somites and ultimately more vertebrae in long-tailed forest compared to short-tailed prairie mice.

Tail length has long been used as an indicator of habitat occupancy, with longer tails associated with arboreality even among closely related species (e.g., squirrels: Hayssen 2008; murine rodents: Nations et al. 2019, field mice: Štěpánková & Vohralík 2008). In deer mice, this correlation was thoroughly investigated by Osgood (1909), who described two distinct ecotypes – forest and prairie forms – based on several morphological traits, with differences in tail length being the most conspicuous. Previous studies suggested an important role for tail use in arboreal locomotion by demonstrating that tail amputation in mice dramatically decreases balance (Horner 1954, Buck et al. 1925, Siegel 1970). Based on these data, a clear hypothesis emerged: naturally evolved tail-length differences in deer mice may be important for performance in arboreal climbing (Horner 1954, Thorington 1970, Kaufman & Kaufman 1992). Recent biomechanical modeling suggests that the longer, heavier tails allow forest deer mice to better control their body roll, as when traversing narrow rods (Hager & Hoekstra 2021). Indeed, in the subspecies we studied here, we found striking differences in a rod-crossing assay – with forest deer mice falling fewer times and completing more crosses than prairie mice – consistent with experimental studies in other populations and species (e.g., Imaizumi 1978, Le Berre & Le Guelte 1993, Layne 1970, Dewsbury et al. 1980). While horizontal climbing on a narrow rod does not capture all the complexities of arboreal locomotion in the wild, deer mice are known to cross narrow twigs in nature (Graves et al. 1988), nor does this assay allow us to disentangle the roles of any behavioral (e.g., balance, skilled movements) or additional morphological differences (e.g., foot size, whisker length) that also may contribute to climbing performance. Nonetheless, these heritable, ecotype-specific differences in rod-crossing ability, in the expected direction, are likely to be at least partly, if not largely, driven by differences in tail morphology.

Genetic mapping allowed us to characterize the genomic architecture underlying total tail length and its constituent components – caudal vertebral length and number – both of which consistently differ between forest and prairie ecotypes across North America (Kingsley et al. 2017). In this species, tail-length differences are largely controlled by six major-effect loci on six different chromosomes. Notably, mapping studies in other wild vertebrates also identified multiple QTL associated with variation in caudal vertebrae (e.g., threespine sticklebacks, Berner et al. 2014, Miller et al. 2014; medaka, Kimura et al. 2012). Because the total variation explained by these six regions together is 24%, this also suggests that many additional loci of small effect were not detectable given the size of our mapping population. Thus, a role for *Hoxd13* would be accompanied by several (possibly many) other genes in establishing vertebra number differences between ecotypes. Of the six loci, three are associated with vertebra length and the other three with vertebra number, consistent with the observation that these traits are not strongly correlated in F2 hybrids. Similarly, artificial selection for increased tail length in replicate lines of laboratory mice resulted in one line with longer vertebrae and the other with more vertebrae (Rutledge et al. 1974). That these traits are genetically separable raises the possibility that the correlation between length and number across the the species could be due to biomechanical constraints (e.g., a trade-off between tail stiffness and flexibility), but modeling does not find support for tail curvature, at least, being strongly influenced by the relative changes in length or number of tail vertebrae in deer mice (Hager & Hoekstra 2021). Instead, the repeated evolution of coincident length and number differences may be due to selection on increased overall tail length by either mechanism when standing variation and/or new mutations are plentiful and exist or appear at roughly the same frequency.

Support for natural selection on tail length in deer mice stems from multiple lines of evidence. First, tail length correlates with habitat, even when corrected for genetic relatedness (Kingsley et al. 2017). Tail-length differences are maintained despite high levels of gene flow connecting forest and prairie populations (Yang & Kenagy 2011). Our QTL mapping results provide additional, independent evidence that supports a possible role of selection: all six detected tail-length QTL have allelic effects in the same direction as the overall tail length difference between ecotypes (i.e., forest alleles are always associated with longer tails and prairie alleles with shorter tails), a result unlikely to occur by chance. Importantly, these findings are all consistent with the hypothesis of divergent selection acting on tail length: that not only are long tails favored in forest habitat, but also short tails are favored in prairie habitat. In the latter case, long tails are likely costly to produce, are a source of heat loss, can be subject to injury, and/or may be an additional target for predation (Thorington 1970, Shargal et al. 1999, Hayssen 2008); therefore, without the benefit of, for example, improving climbing performance, the cost of having a long tail outweighs the benefit in terrestrial mice inhabiting open, prairie habitats.

Our mapping study also allowed us to narrow in on promising candidate genes contained within the QTL intervals. We found that two of the three QTL influencing vertebra number contain Hox gene clusters: Hoxa and Hoxd. While our approach does not allow us to rule out the involvement of other genes in these intervals, Hox genes are especially intriguing candidates in light of recent studies: in addition to specifying tail identity, the Hox13 paralogs also have been proposed to control axis termination (Diaz-Cuadros et al. 2021). First, the most 5’ Hox genes, those of the paralogy group 13, are expressed at the tip of the elongating embryonic tail (Deschamps & Duboule 2017), where these genes are known to terminate axial elongation by inhibiting the effects of more anterior Hox genes and repressing Wnt activity (Denans et al. 2015, Beccari et al. 2016, Sheth et al. 2016). For example, in *Mus*, loss of *Hoxb13* leads to the formation of extranumerary caudal vertebrae (Economides et al. 2003), while its overexpression results in premature truncation of the tail (Young et al. 2009). Consistent with these studies, we found lower levels of *Hoxd13* in long-tailed forest mice compared to higher levels in short-tailed prairie mice. Studies that manipulated the expression of *Hoxd13* alone in *Mus* have not detected changes in tail length (e.g., Tschopp et al. 2009), but this is consistent with the subtle phenotypic effect of the QTL containing *Hoxd13* in deer mice (PVE = 3.1%). Moreover, equivalent changes in *Hoxd13* may also have different effects or effect sizes in species other than *Mus*. Thus, together, these genetic data indicate that *Hoxd13* is a strong candidate for contributing to the evolution of caudal vertebra number differences in deer mice: *Hoxd13* is expressed in an appropriate time and place (i.e., in the developing tail bud during somitogenesis); it is regulated by *cis*-acting variants as expected for a causative locus identified via QTL mapping; and the direction of expression difference between ecotypes is consistent with its known function..

To better understand the possible role of *Hoxd13* during development, we explored the developmental mechanisms that ultimately lead to differences in vertebra number. Evolution of vertebra number is likely to require changes to the parameters of axial segmentation and/or elongation. Previous studies investigating somite number differences in snakes *vs*. non-snakes (Gomez et al. 2008), between inbred lines of medaka (Kimura et al. 2012), and zebrafish *hes6* and *hes7* timing mutants (Schröter & Oates 2010, Harima et al. 2013) implicated changes to the rate of segmentation: faster somite formation rates produce more, smaller somites. By contrast, here we did not find evidence that forest mice have smaller somites or that somites form at a faster rate, but instead found that forest deer mice develop a larger amount of PSM tissue in the post-hindlimb region, suggesting a faster rate of axis elongation given the same rate of somite formation. Because somitogenesis is thought to end when posterior elongation ceases and the somite formation front “catches up” to the tip of the tail, a longer post-hindlimb PSM is predicted to result in more somites (Gomez & Pourquie 2009, Kimelman 2016).

Recent work on axial development has shown how Hox expression can influence, in addition to vertebral identity, the overall length of the vertebral column by regulating posterior axial extension (Young et al. 2009, Aires et al. 2019, Robinton et al. 2019). This effect is mediated by regulation of progenitor cells, NMPs, that give rise to the PSM; indeed, a single-cell RNA-seq study in *Mus* found that *Hoxd13* is expressed in NMPs (Guillot et al. 2021). In both mice and fish, posterior Hox genes, especially the Hox13 paralogs, act to maintain this progenitor population, at least in part by inhibiting Wnt and FGF signaling (Denans et al. 2015, Aires et al. 2019, but see Ye & Kimelman 2020). Indeed, one prediction of a reduction in *Hoxd13* expression is an increase in Wnt signaling that would sustain the NMP population (Denans et al. 2015). Our data show that *T* and *Cdx2*, which are Wnt targets (Chawengsaksophak et al. 2004, Martin & Kimelman 2008, van de Ven et al. 2011), have higher expression in forest versus prairie mice. Because *T* is essential for production of PSM, this provides a potential mechanism by which decreased *Hoxd13* expression in the forest tail bud could result in an elongated embryonic axis. However, not all Wnt target genes (e.g., *Lef1, Axin2*) show a similar pattern. Thus, the precise details of how changes in *Hoxd13* expression may promote or maintain the larger NMP population have yet to be fully explained. Nonetheless, these results suggest that differences in the size of the axial progenitor pool, likely influenced by *Hoxd13* expression differences, underlie differences in PSM size and thus vertebra number in deer mice.

Developmental geneticists have long known that mutations in Hox genes can affect segmental identity in bilaterian animals (Lewis 1978, Akam 1989), although these lab-derived homeotic “monsters” are clearly less fit than the wild type. Nonetheless, this potential, along with the correlation of Hox expression patterns with body segments, led many to enthusiastically hypothesize that changes in Hox genes could underlie major morphological shifts in animal body plans in nature (Gaunt 1994, Burke et al. 1995, Averof & Patel 1997, Lemons & McGinnis 2006). Thus, while HOX protein sequences are conserved due to their pleiotropic roles in development (McGinnis et al. 1984, Krumlauf 1994, Carroll 1995), it was unclear if regulatory changes at Hox loci contribute to segmental evolution in natural populations, especially in vertebrates. Here, we provide evidence that *cis*-acting mutation(s) causing gene expression change in *Hoxd13*, through developmental changes to the PSM and its progenitor cells, contributes to segment number variation within a single species of deer mouse. Together this work, and the parallel work showing *cis*-regulatory changes in *Hoxdb* associated with spine number variation in stickleback fish (Wucherpfennig et al. *in review*), demonstrate how Hox genes can contribute to adaptive morphological evolution even on microevolutionary scales in the wild.

## ACKNOWLEDGEMENTS

We thank Terry Capellini, Denis Duboule, Shuonan He, David Kingsley, Olivier Pourquié, Cliff Tabin, Julia Wucherpfennig and members of the Pourquié lab for providing helpful feedback about data and/or the manuscript; Terry Capellini, Pushpanathan Muthuirulan, Mariel Young, Phil Grayson, and Jenny Chen for advice on molecular and computational methods; Mark Omura for shipping samples; Caroline Hu for collecting embryos; and Emily Jacobs-Palmer for help in the field and in the lab. The Bauer Core Facility at Harvard University supplied library prep and sequencing services. The computations in this paper were run on the FASRC Odyssey and Cannon clusters supported by the FAS Division of Science Research Computing Group at Harvard University. This work was supported in part by Putnam Expedition Grants from the Museum of Comparative Zoology to EPK and the Robert A. Chapman Memorial Scholarship for the study of Vertebrate Locomotion to EPK and ERH. ERH was supported by an NIH Training Grant to Harvard’s Molecules, Cells, and Organisms graduate program (NIH NIGMS T32GM007598) and by the Theodore H. Ashford Fellowship. OSH was supported by a National Science Foundation Graduate Research Fellowship, a Harvard Quantitative Biology Student Fellowship (DMS 1764269), and the Molecular Biophysics Training Grant (NIH NIGMS T32GM008313). JML received support from the European Molecular Biology Organization (ALTF 379-2011), the Human Frontiers Science Program (LT001086/2012), and the Belgian American Educational Foundation. HEH is an Investigator of the Howard Hughes Medical Institute.

## AUTHOR CONTRIBUTIONS

EPK and HEH conceived experiments. EPK performed the genetic cross, phenotyping and related analyses, embryo measurements and analysis, ecotype RNA-seq and F1 RNA-seq analyses, and CRISPR experimental design and analysis. ERH designed, conducted and analyzed behavioral assays. JML called variants for cross genotypes, generated personalized genome assemblies and annotations, and provided advice for RNA-seq analyses. KMT and OSH made RNA-seq libraries. CK and BIN managed CRISPR mice and genotyped and collected *Mus* pups. BIN collected tissues for RNA-seq. EPK, ERH, OSH, JML, and HEH wrote the manuscript with input from all authors.

## METHODS

### Animals

We focused on two subspecies of deer mice, *Peromyscus maniculatus*, representing the forest (*P. m. nubiterrae*) and prairie (*P. m. bairdii*) ecotypes. Forest mice were descendants of 16 wild-caught deer mice that we captured from maple-birch forest in Westmoreland County, Pennsylvania in 2010 (described in Kingsley et al. 2017). Prairie mice were descendants of mice obtained from the Peromyscus Genetic Stock Center (University of South Carolina), originally captured in Washtenaw County, Michigan in 1948.

Mice were housed at 23°C on a 16:8-hour light:dark cycle in standard mouse cages (Allentown Inc.) with corncob bedding (The Andersons, Inc.), cotton nestlet (Ancare), Enviro-Dri (Shepherd Specialty Papers), and either a red tube or a red hut (BioServ). Mice were housed in same-sex groups of two to five individuals and provided with water and mouse chow (LabDiet Prolab Isopro RMH 3000 5P75) *ad libitum*. All breeding colonies and experiments were conducted under and approved by the Harvard IACUC protocol 11-05.

### Behavioral assay

To measure an ecologically-relevant aspect of climbing performance in which the tail may play a role, we designed a rod-crossing assay, similar to that used by Horner (1954). In brief, we built a custom arena consisting of two 32 cm x 14.5 cm white acrylic platforms (McMaster-Carr), elevated 65 cm above the floor and connected by a 44 cm long, 5/32” (0.4 cm) diameter stainless steel rod (Fig. 2A). To start each trial, we placed a naive, adult mouse on the platform for a brief 1-min habituation and then allowed the mouse to voluntarily explore the arena. Trials lasted 5 mins after the start of the first cross (defined as when the mouse first placed all four feet on the rod) or for a maximum of 10 mins if the mouse never initiated rod crossing. We filmed the trials at 240 fps, 720 × 1280 pixel resolution, using two GoPro Hero 4 Black cameras mounted on tripods (one top view and one side view). We performed all assays during the light phase, between zeitgeber time (ZT) 10 and 14 (with ZT 0 defined as lights-on). Between trials, we cleaned the arena with 70% ethanol and allowed it to dry fully. Each mouse was tested once, between 55 and 70 days of age.

For each trial, we manually scored behaviors, including crossing the rod and falling. Specifically, we defined a ‘cross’ as the time between a mouse placing all four feet on the rod and when the last foot was removed from the rod. For each cross, we scored whether the mouse fell (i.e., lost all contact with the rod before remounting the platform). In cases in which the mouse did not fall, we noted whether the mouse completed the cross by reaching the other platform (i.e., whether the mouse touched the opposite platform at any point during the trial). We report results for all mice that climbed onto the rod at least once during a trial (forest, n = 32 of 35 complete trials; prairie, 31 of 46 complete trials).

If a mouse fell or jumped from either the rod or the platform, the experimenter stopped the assay and replaced the mouse on the platform. If a mouse jumped from the platform more than five times during a trial, the trial was discontinued and not analyzed further (forest, n = 5; prairie, n = 5).

### Statistical analysis of behavior

We analyzed behavior data using generalized linear mixed models (family = “binomial”, *lme4* package v. 1.1, in R v. 3.6.2 [Bates et al. 2015, R Core Team 2021]), including data for only the first eight cross attempts, as no prairie mice crossed more than eight times during the trial while forest mice crossed up to 34 times. Each response variable was binary (“fell” or “completed”). We fit models with the following sets of fixed effects: cross index alone (null model; cross number is included to account for possible effects of experience); ecotype alone (i.e., no effect of experience); additive effects of ecotype and cross; or an interaction between ecotype and cross (i.e., different effects of experience in the two ecotypes). Each model also included *individual* as a random effect. We compared these models using likelihood ratio tests (implemented in the anova function, “stats” package).

### Genetic cross

#### Forest-prairie F2 hybrid intercross

To produce a genetic mapping population, we established a reciprocal intercross between two ecotypes: forest (*P. m. nubiterrae*) and prairie (*P. m. bairdii*). The mapping cross consisted of two families, each founded by two animals: family “0”: female *bairdii* x male *nubiterrae*; family “1”: female *nubiterrae* x male *bairdii*. Cross parents were siblings. We established 14 F1 breeding pairs, which when intercrossed produced 495 F2 hybrids (family 0, n = 211; family 1, n = 284) for analysis. F2 hybrids were sacrificed between ages 70–300 days and were measured for gross morphology (total length, tail length, and body mass).

### Skeletal measurements

We measured lengths of limb- and tail-bones in 12 forest and 12 prairie, 14 F1 hybrid animals, and 495 F2 hybrid animals from x-ray radiographs. We used a digital x-ray system (Varian Medical Systems, Inc.) in the Harvard Museum of Comparative Zoology Digital Imaging Facility to obtain radiographs of whole specimens mounted such that the plane containing the anterio-posterior and medio-lateral axes was parallel to the imaging plane. We measured all traits with Fiji/ImageJ (Schindelin et al. 2012); we included a standard to determine scale. In total, we measured up to 32 sacral and caudal vertebrae, maximum caudal vertebra length and caudal vertebra number, as well as total sacrum length and total tail length (Fig. S1).

Most bone-length traits were correlated with body size in our cross, therefore we corrected for body size using linear regression on sacrum length. Sacrum length, a section of the vertebral column that is anterior to the caudal vertebrae and does not significantly differ in length between ecotypes (Wilcoxon test, W = 38, *p* = 0.1), represents a standard for body size (sacrum length vs. body mass: in F2s, Pearson’s *r =* 0.55, 95% CI: 0.48–0.60; vs. ruler-measured body length: *r* = 0.62, 95% CI: 0.56–0.67). We corrected for body size by regressing raw trait measures against the sum length of the six sacral vertebrae and added the residuals from that regression to the trait mean to align the corrected measurements in the ranges of the raw measurements.

To describe variation in tail length, we used three summary statistics: (1) the number of caudal vertebrae (all vertebrae posterior to the six sacral vertebrae), (2) the length of the longest vertebra in the tail, and (3) the total length of the tail, measured from the x-ray radiographs (Fig 3A). We explored the pairwise correlations among traits in the F2 animals and conducted a PCA (as implemented in the “principal” function in the *psych* package in R; [Revelle 2021; R Core Team 2021) using measurements with the standard deviations for each trait scaled to 1 and centered the means of each trait to 0. The first three components account for 78% of the variance in sacral and caudal vertebral lengths.

### Genotyping and linkage map construction

We genotyped parent, F1, and F2 animals using double digest restriction-site associated DNA sequencing (ddRADseq; Peterson et al. 2012). Briefly, we extracted genomic DNA from alcohol-preserved liver tissue with the AutoGenprep 965 (AutoGen; Holliston, MA), digested it with EcoRI and MspI (New England Biolabs; Ipswich, MA) and ligated end-specific adapters, P1 and P2 that include individual barcodes and biotin labels, respectively. Next, we combined samples into 48-individual pools and size-selected each pool to 216–276 bp using a Pippin Prep (Sage Science; Beverly, MA), after which we used streptavidin beads (Dynabeads M-270, Life Technologies; Carlsbad, CA) to eliminate fragments without P2 adapters. We PCR-amplified these pools (10 cycles) with an indexed primer. Using a TapeStation (Agilent; Santa Clara, CA), we quantified the mass of these pools (ranged from 0.7 to 5.0 nM) and combined them in equimolar ratios. Finally, we sequenced these pools in 150-bp paired-end rapid runs on an Illumina HiSeq 2500 to ∼600K reads per sample.

We processed the sequence reads using custom Python software (described in Peterson et al. 2012; github.com/brantp/rtd). In brief, this software used Stampy to map merged paired-end reads to the *P. maniculatus* genome scaffolds (GCA_000500345.1) and then combined reads by individual into BAM files with Picard (broadinstitute.github.io/picard/). We then used GATK (McKenna et al. 2010, DePristo et al. 2011) to call variants with UnifiedGenotyper. From 4.3e8 raw reads, this analysis produced 1.1e7 called variants. We hard filtered these variants for those that were fixed between the prairie and forest parents of the cross, those with QD > 5, GQ > 30, and those present in more than half the F2 individuals (using HTSeq; Anders et al. 2015). This filtering produced 4,296 variants, which we used to construct a linkage map using R/qtl, closely following the procedure outlined by Broman & Sen (2009). The resulting map had 24 linkage groups, corresponding to the haploid number of chromosomes in *P. maniculatus* (Singh & McMillan 1966), comprised of 2,618 markers with an average spacing between markers of 0.7 cM, and a maximum spacing of 23.1 cM.

### Quantitative trait locus (QTL) mapping

We used R/qtl (Broman & Sen 2009) to identify regions of the genome in which genetic variation was statistically associated with variation in skeletal traits. For all bone-length traits, we performed standard interval mapping with the extended Haley-Knott method (“ehk” in the R/qtl *scanone* function) including *sex, age*, and *sacrum length* as additive covariates. Because the number of caudal vertebrae (*count*) was not continuous and not normally distributed (Shapiro-Wilk test: W = 0.90, p < 1e-15), we used the nonparametric method for mapping. We used permutation tests (n = 1000 permutations for autosomes, n = 26,312 for the X chromosome) to determine significance thresholds for each trait (Churchill & Doerge 1996).

To assess the effect sizes of each QTL and the amount of variance each locus explained, we used multiple-QTL models and drop-one analysis in R/qtl. Using the p < 0.05 significance thresholds as determined by permutation tests, we fit models for each trait with the genotypes at markers with the highest LOD scores in each significant QTL as explanatory variables as implemented in *fitqtl*. The models for length traits include *sex, age*, and *sacrum length* as additive covariates.

We assessed evidence for selection on tail length using the direction of QTL effects with the QTLSTEE (Orr 1998) and with the ratio of parental and F2 variances with the *v* test (Fraser 2020). For the *v* test, we used a conservative assumption of additivity (c = 2) and estimated H^2^ using parental, F1, and F2 variances (Lynch & Walsh 1998).

### Embryo collection

We generated embryos of approximate ages (E11.5–E15.5) from each ecotype. Because *Peromyscus* mice experience postpartum estrus (Dewsbury 1979), we set the date of conception as the birth date of a female’s last litter and then confirmed these ages using a developmental time series of *Peromyscus* (Manceau et al. 2011, Davis & Keisler 2016).

### RNA-seq of embryonic tail tissue

We dissected post-anal tail tissue from 35 embryos (forest, n = 18; prairie, n = 17) at Theiler Stages 15–20 (E12.5–E15.5), timepoints relevant to tail somitogenesis (Theiler 1989). We extracted total RNA using the PicoPure RNA Isolation kit (ThermoFisher Scientific) and constructed RNA-seq libraries using PrepX poly-A and library prep kits on an Apollo 324 System, following the manufacturer protocol. We sequenced libraries on two lanes of 150-bp paired-end runs on an Illumina HiSeq 2500 to ∼30 million reads/sample.

To measure allelic expression bias in F1 hybrid embryos, we dissected embryonic tails at E12.5 (n = 4) and E14.5 (n = 4) and extracted RNA using 50 µl Direct-zol (Zymo Research) following the manufacturer’s protocol and used the same library preparation procedures as for the parental samples. We sequenced libraries on one 150-bp paired-end run on an Illumina NovaSeq SP flowcell to ∼45 million reads/sample.

We assessed differential expression using an established workflow, following Bendesky et al. (2017). Briefly, we trimmed reads using Cutadapt (Martin 2011) via Trim Galore! (github.com/FelixKrueger/TrimGalore) and mapped reads to the *P. maniculatus* genome (Pman2.1.3; GCA_003704035.3) (forest and prairie libraries) or a custom hybrid genome created from variants called from RNA-seq reads (F1 libraries) using STAR aligner (Dobin et al. 2013). Eighty-five percent of annotated transcripts in the hybrid genome have at least 1 variant that allowed allele assignment, including our top five candidate genes. We quantified transcripts using RSEM (Li & Dewey 2011) and used edgeR (Robinson et al. 2010) and limma-voom (Law et al. 2014) to compare transcript abundance between ecotypes at both early (E12.5–13.5) and late (E14.5–15.5) stages. We normalized libraries using the TMM method, as implemented in edgeR, and ranked differentially expressed genes by the empirical Bayes (eBayes) method in limma.

### Identification of candidate genes

To prioritize candidate genes related to skeletal variation within QTL intervals, we first calculated 95% confidence intervals for each QTL using the *bayesint* function in R/qtl. We extracted names of genes in the QTL intervals from the *P. maniculatus* (Baylor 2013) genome annotation and used the resulting list of gene names to cross-reference with alleles from the Mouse Genome Informatics (MGI) Mammalian Phenotype Browser (www.informatics.jax.org/searches/MP_form.shtml) that have “limb/digits/tail” phenotypes (Table S3).

### CRISPR-HDR for HOXD13 amino acid mutation

To test the effect of *Hoxd13* amino acid mutations on tail development, we conducted a CRISPR-Cas9 homology-directed repair experiment in *Mus*. Specifically, we designed a guide RNA and homology-directed repair (HDR) template to insert a single alanine into the *Mus Hoxd13* locus at amino acid position 109 (*Hoxd13*^A109^). The sequences of the synthesized guide RNA (Synthego) and single-stranded HDR template (IDT) are provided in Table S4. These were injected along with Cas9 protein (IDT) into C57BL/6J zygotes by the Harvard Genome Modification Facility.

We amplified and sequenced the edited allele (primer sequences in Table S4) from tail-tip DNA and assessed editing efficiency using the Synthego ICE tool (ice.synthego.com). We mated the three males and three females with the highest editing efficiency to wild-type animals and then intercrossed siblings to produce F2 offspring (+/+, n = 22; +/d13^A109^, n = 55; d13^A109^/d13^A109^, n = 37). A successful edit destroys a PstI restriction site, so we genotyped P0 F2s using the same primers followed by PstI restriction digestion of the resulting amplicon. To confirm that the correct edit was made, we sequenced *Hoxd13* exon 1 in a subset of F2 animals (n = 4 homozygotes for each allele from each family, 24 total); we did not find any off-target mutations in these sequences.

#### Postnatal vertebral counts

We used whole-mount bone/cartilage staining to compare caudal vertebra counts in laboratory-reared neonatal (P0) pups of forest (n = 6) and prairie (n = 6) ecotypes, and of +/+ and *Hoxd13*^A109^*/Hoxd13*^A109^ CRISPR-HDR (n = 114) *Mus* F2s. We stained bone and cartilage with alizarin/alcian following Rigueur & Lyons (2014) and counted all recognizable segments in the tail, including non-ossified cartilage condensations at the caudal tip (Fig. 5C; Fig. S5A).

Investigators were blind to ecotype/genotype when counting segments.

### Measurement of PSM and somite lengths

To compare tissue dimensions in fixed embryos, we sacrificed females and dissected embryos in PBS, then fixed the embryos in phosphate-buffered 4% formaldehyde for 14–24 hours at 4°C. We stained whole embryos with 1µg/mL DAPI for 30 minutes and photographed them with a Zeiss mRc camera on a Zeiss steREO Discovery V.12 dissecting microscope that was scale calibrated. We used the linear measurement tool in Fiji/ImageJ (Schindelin et al. 2012) to measure somite and PSM lengths. We analyzed these data in R (R Core Team 2021) and made plots using *ggplot2* (Wickham 2009).

### Embryonic tail explant culture and time lapse imaging

To obtain precise measurements of segmentation and axial extension parameters, we cultured posterior embryonic tissues and time-lapse imaged them. We dissected E12.5–E15.5 embryos in DMEM that was pre-warmed to 37°C, dissected the portion of the embryo caudal to the hind limb bud, and transferred that explant to an uncoated Mat-Tek glass-bottomed culture dish also containing prewarmed DMEM. We then transferred the dish containing explant to a culture chamber at 37°C with a humidified carbon dioxide (5%) line on a Zeiss Cell Observer (Harvard Center for Biological Imaging). We used Zen 2012 (Zeiss) software to take images every ten minutes over a 12–14 hour period while the explant formed somites and underwent axial extension. We took a Z-stack for each time point and used the “Extended Depth of Focus” function in Zen to collapse the stack into a single image for each time point. From these time-lapse movies, we obtained basic information about the timing of segment formation using Fiji/ImageJ to mark the formation of somite boundaries on individual frames. All explants settled slightly during the first 90–120 minutes; for all time-lapse movies, we discarded the first 12 frames.

### Immunostaining and cell counting

We dissected embryos from pregnant female forest and prairie mice (forest, n = 6; prairie, n = 5), and fixed embryos in phosphate-buffered 4% formaldehyde for 14–24 hours at 4°C. We rinsed with PBS, then embryos were graded through 10% sucrose/PBS (1 hour at 20°C), 30% sucrose/PBS (overnight at 4°C), and then mounted in OCT medium and frozen. We cryosectioned tails in the sagittal plane at 14 µm per section, then immunostained with anti-Sox2 (R&D Systems MAB2018; 1:500), anti-Brachyury/T (R&D Systems AF2085; 1:500) and fluorophore-conjugated secondary antibodies (anti-mouse-AlexaFluor555 and anti-goat-AlexaFluor488; 1:500; ThermoFisher), each overnight at 4°C. We counterstained with 1µg/mL DAPI (30 min at 20°C) and imaged sections with a Zeiss LSM710 confocal microscope with a Plan Apo 20x/0.8 Air DIC II objective. We outlined regions of tail bud mesenchyme (Fig. 6E) in a single section per embryo closest to the midline and counted by hand the total number of DAPI-labeled nuclei, SOX2-positive cells, T-positive cells, and SOX2/T co-labeled cells in this region. Investigators were blind to ecotype when counting cells.

### Data and code availability

The raw and processed forest, prairie, and F1 RNA-seq data have been uploaded to NCBI GEO and awaiting accession numbers. QTL mapping files and code will be available on Dryad. All other data and materials will be made available upon reasonable request.

## SUPPLEMENTAL INFORMATION

**Figure S1.**
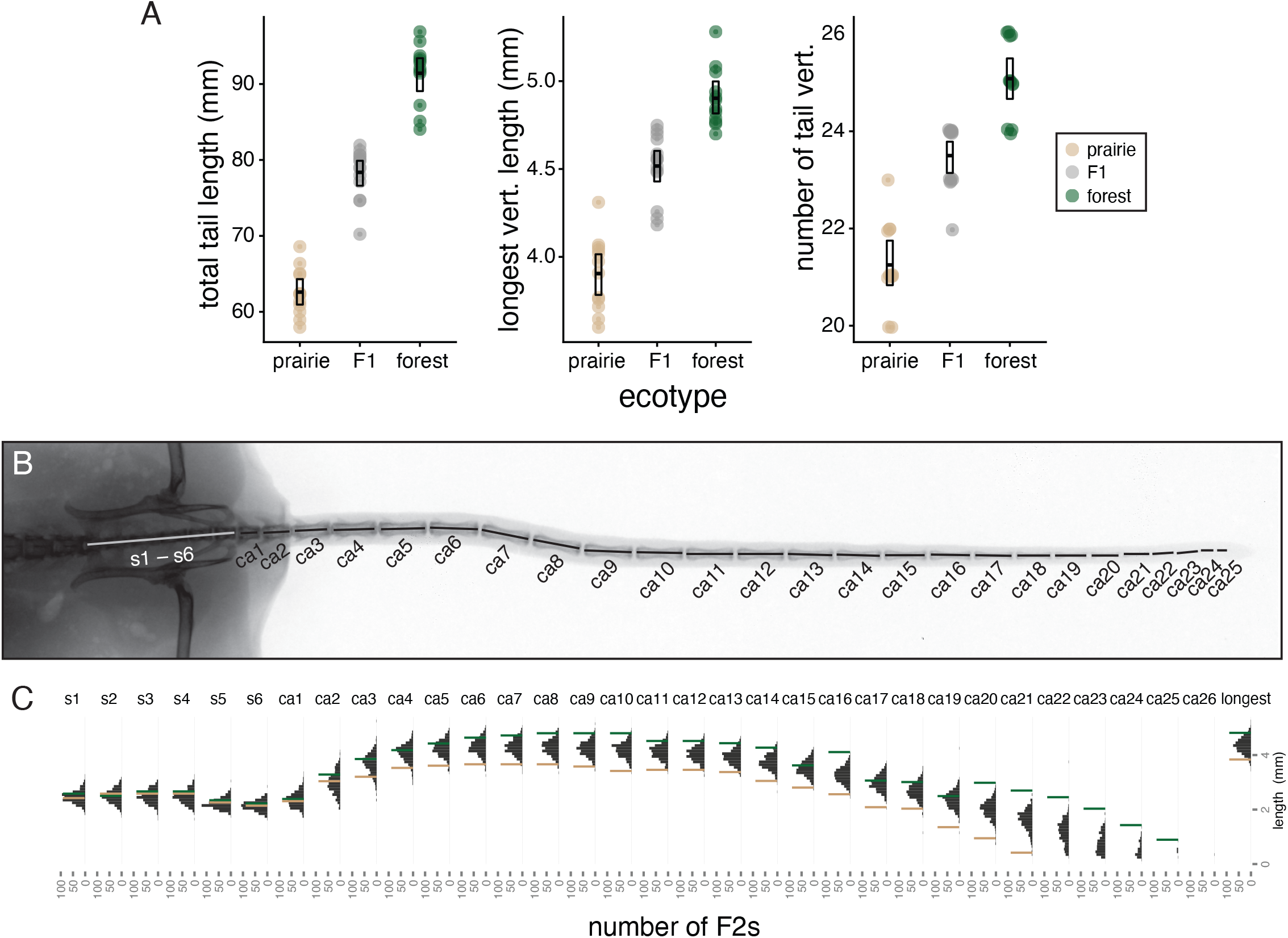
Skeletal trait measurements in parental and hybrid mice. (Related to Figs. 1 & 3) **A**. Total tail length, length of longest vertebra, and number of tail vertebrae in lab-reared prairie (n = 12, tan), F1 (n = 14, gray), and forest (n = 12, green) mice. Boxes show mean and bootstrapped 95% confidence limits of the mean. **B**. Example of x-ray image used to measure vertebra lengths for sacral (s) and caudal (ca) vertebrae. **C**. Distributions of vertebrae length (for each vertebra, s1-6, ca1-26) in F2 hybrid mice (n = 495). Parental mean values are indicated by lines: forest (green) and prairie (tan).

**Figure S2.**
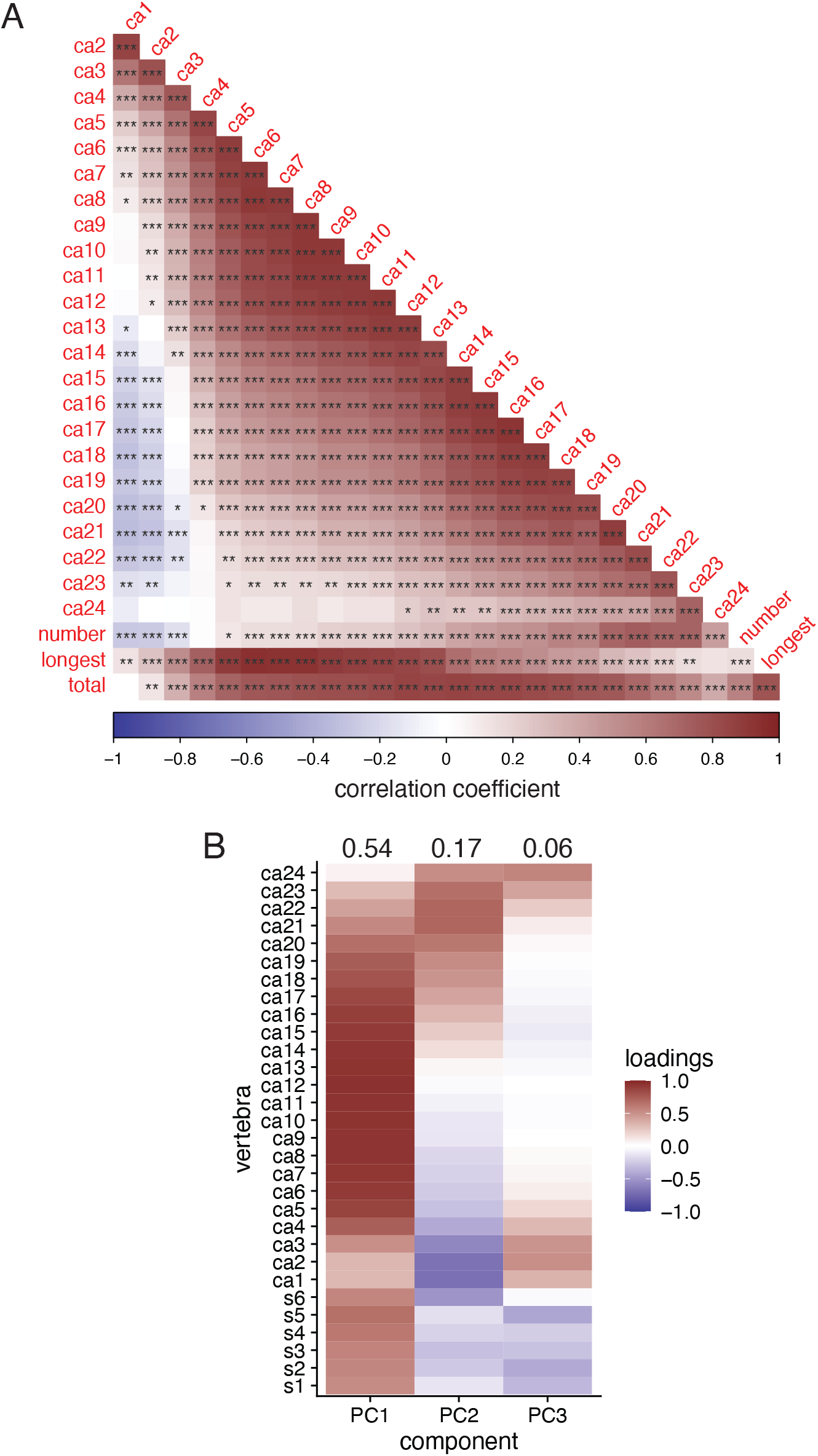
Correlations among tail traits in F2 hybrid mice. (Related to Figs. 1 & 3) **A**. Heatmap of pairwise Pearson correlation coefficients between pairwise tail traits in all F2s (n = 495). All traits except vertebra number are corrected for sacral length (see Methods). **B**. Heatmap of vertebra length loadings from PCA showing that vertebra length traits in different tail regions load onto the first three PCs. Percent variance accounted for by each PC indicated above each column. Abbreviations: s, sacral vertebra; ca, caudal vertebra. * = p < 0.05, ** = p < 0.01, *** = p < 0.001.

**Figure S3.**
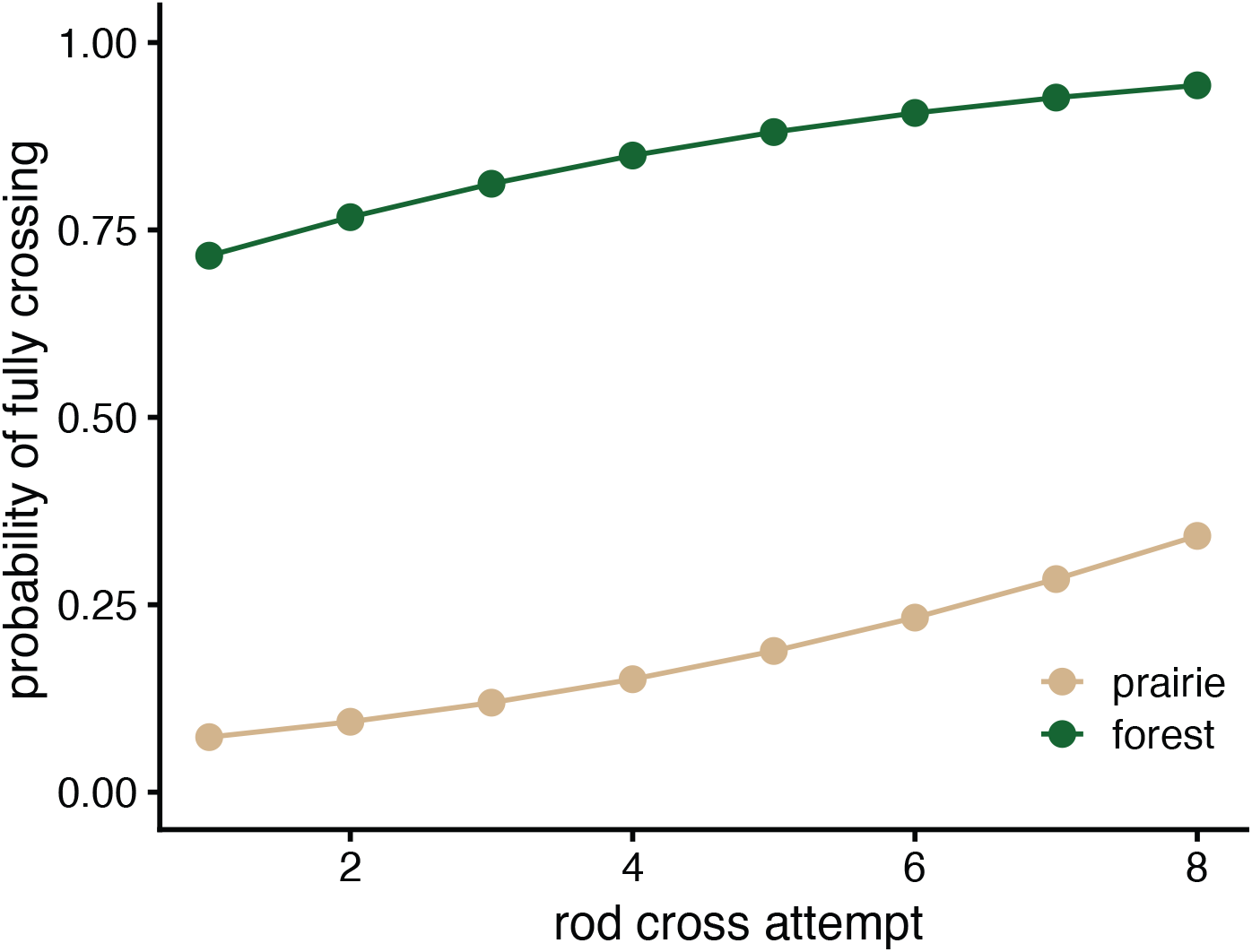
Probability of mice completing a rod crossing. (Related to Fig. 2) The baseline odds (i.e., on the first cross attempt) of a prairie mouse (tan) crossing fully are 0.08:1 (probability 7.3%) and for a forest mouse (green), 2.51:1 (probability: 72%) (p = 8e-8 for effect of ecotype). For each additional cross, the log-odds of a full cross increase by 0.27 (p = 2e-3) for both ecotypes. (Logistic mixed effects model; formula: *complete_cross ∼ ecotype + cross*).

**Figure S4.**
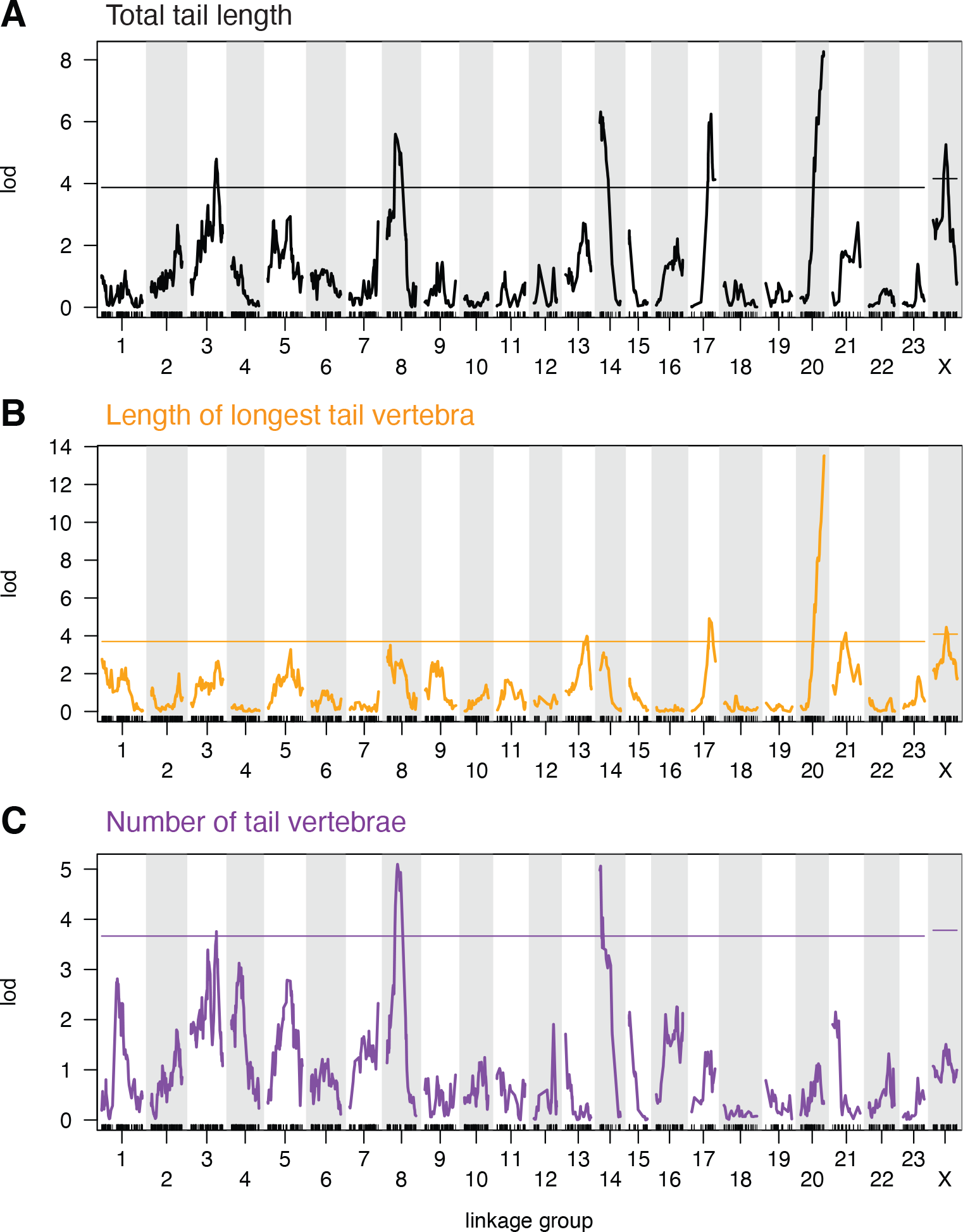
Genome-wide QTL maps for three focal tail traits. (Related to Fig. 3) Statistical association (LOD, or log of the odds, score) between genotype and phenotype across all linkage groups for **A**. total tail length (black), **B**. length of longest caudal vertebra (orange), and **C**. caudal vertebrae number (purple). Horizontal lines indicate genome-wide significance thresholds (p = 0.05) as determined by permutation tests (see Methods). For details of each QTL, see Table S1.

**Figure S5.**
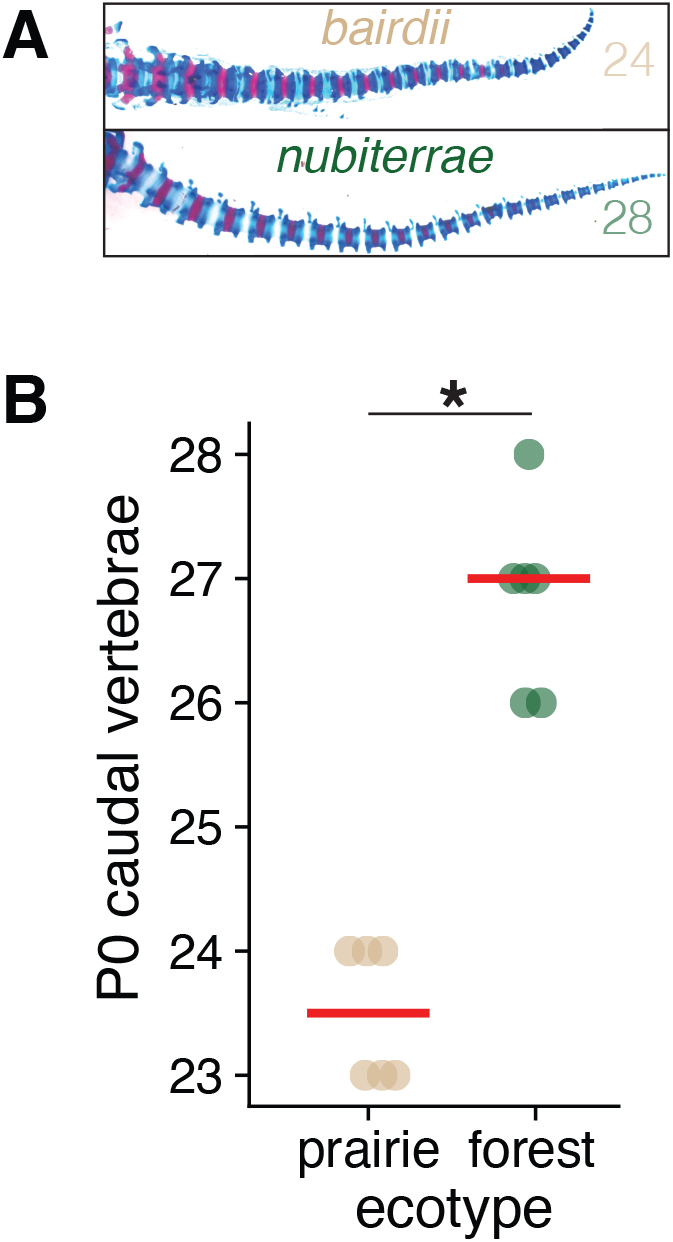
Difference in vertebra number between forest and prairie mice is present at birth. **A**. Examples of postnatal day 0 (P0) tail skeletons stained with alizarin/alcian with caudal vertebra number given. **B**. Caudal vertebra number for P0 forest (green) and prairie (tan) mice (n = 6 for each ecotype). Red lines indicate means. Vertebra number differs significantly (t-test; p = 4e-3).

**Figure S6.**
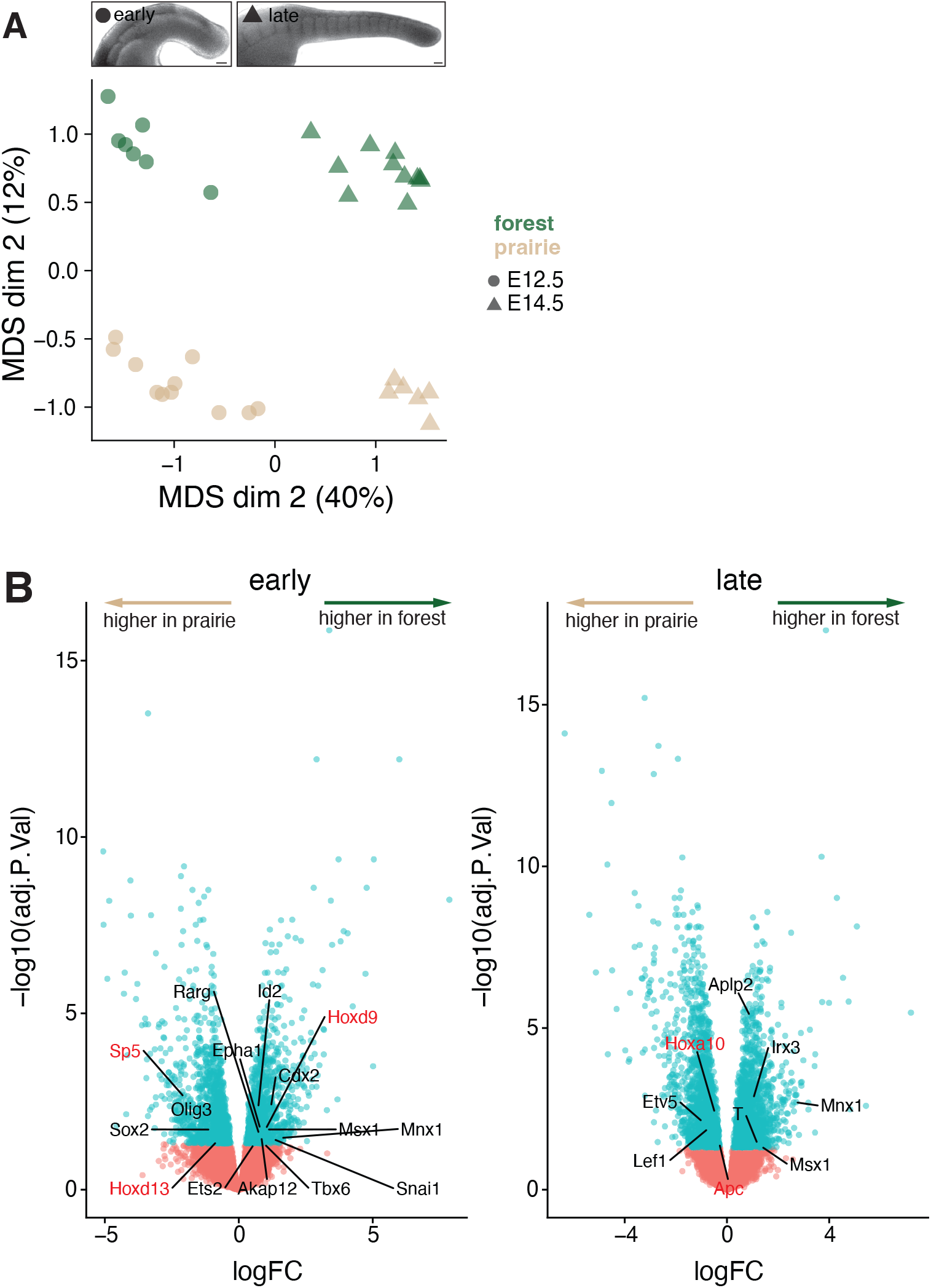
RNA-seq-estimated differential expression between forest and prairie embryonic tail tissues. (Related to Figs. 4 & 6) **A**. Multidimensional scaling (MDS) plot of bulk RNA-seq libraries (forest, green, n = 18; prairie, tan, n = 17) based on top 500 most highly-expressed genes in embryonic tails from early (E12.5–13.5, circle) and late (E14.5–15.5, triangle) embryonic stages of tail elongation. Top panel: Examples of early- and late-stage embryonic tails, tailbud on the right. Scale bar = 100 µm. **B**. Volcano plots showing differentially expressed genes between forest and prairie samples at early and late stages of tail elongation. Blue dots indicate adjusted p-value < 0.05. Gene name labels indicate differentially expressed genes associated with NMPs (black text) or candidates from QTL intervals (red text). logFC = log2 fold change, forest - prairie.

**Figure S7.**
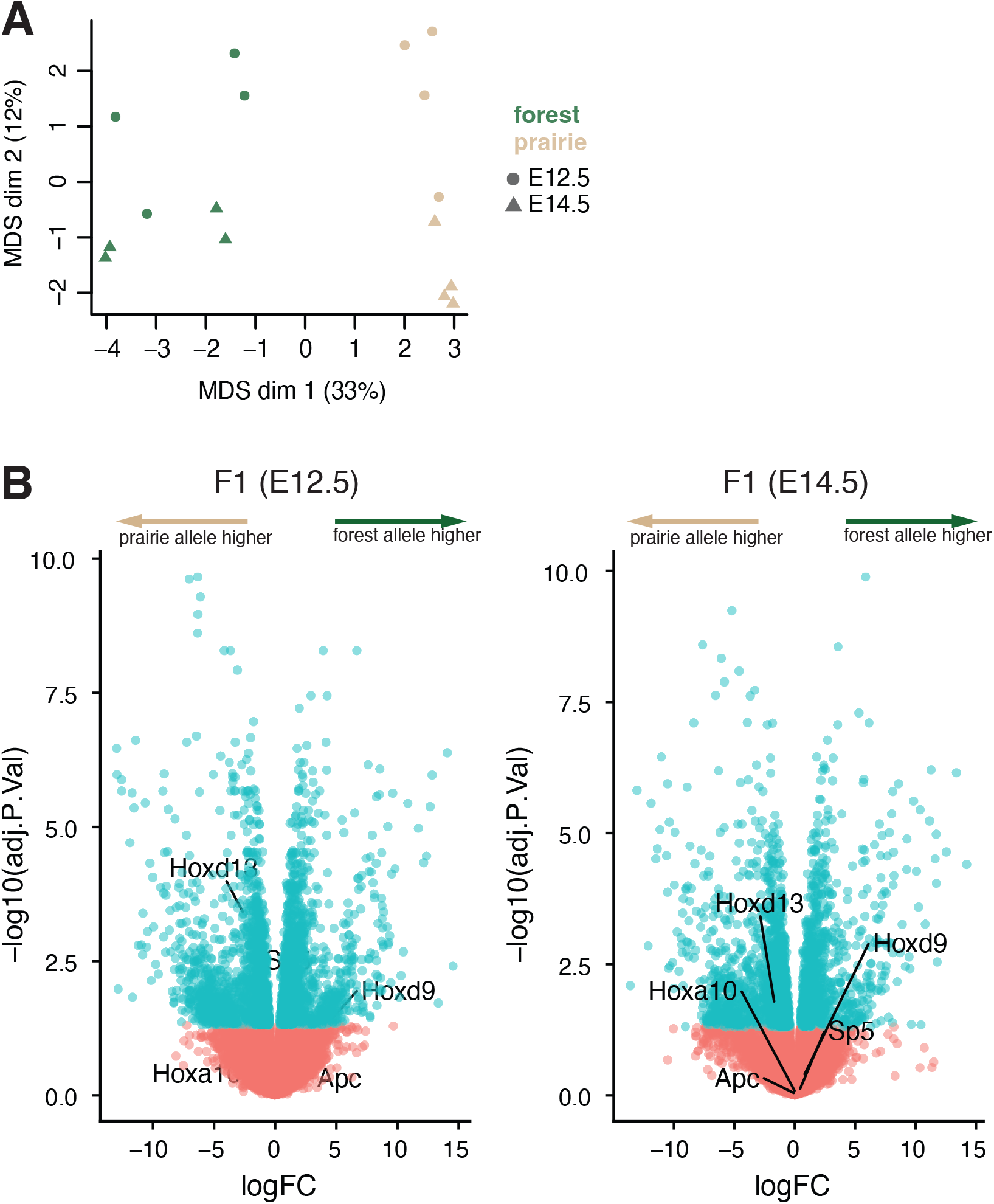
RNA-seq-estimated allele-specific differential expression in forest-prairie F1 hybrids. (Related to Fig. 4) **A**. Multidimensional scaling (MDS) plot of bulk RNA-seq F1 forest-prairie hybrid libraries based on the top 500 most highly-expressed genes in embryonic tails showing forest (green) and prairie (tan) alleles from early (E12.5, circle) and late (E14.5, triangle) embryonic stages of tail elongation. **B**. Volcano plots showing genes with allele-specific expression at E12.5 and E14.5. Blue dots indicate genes with significant allele-specific expression (adjusted p-value < 0.05). Gene name labels show genes that both are in QTL intervals and have MGI tail phenotypes. logFC = log2 fold change, forest allele - prairie allele.

**Figure S8.**
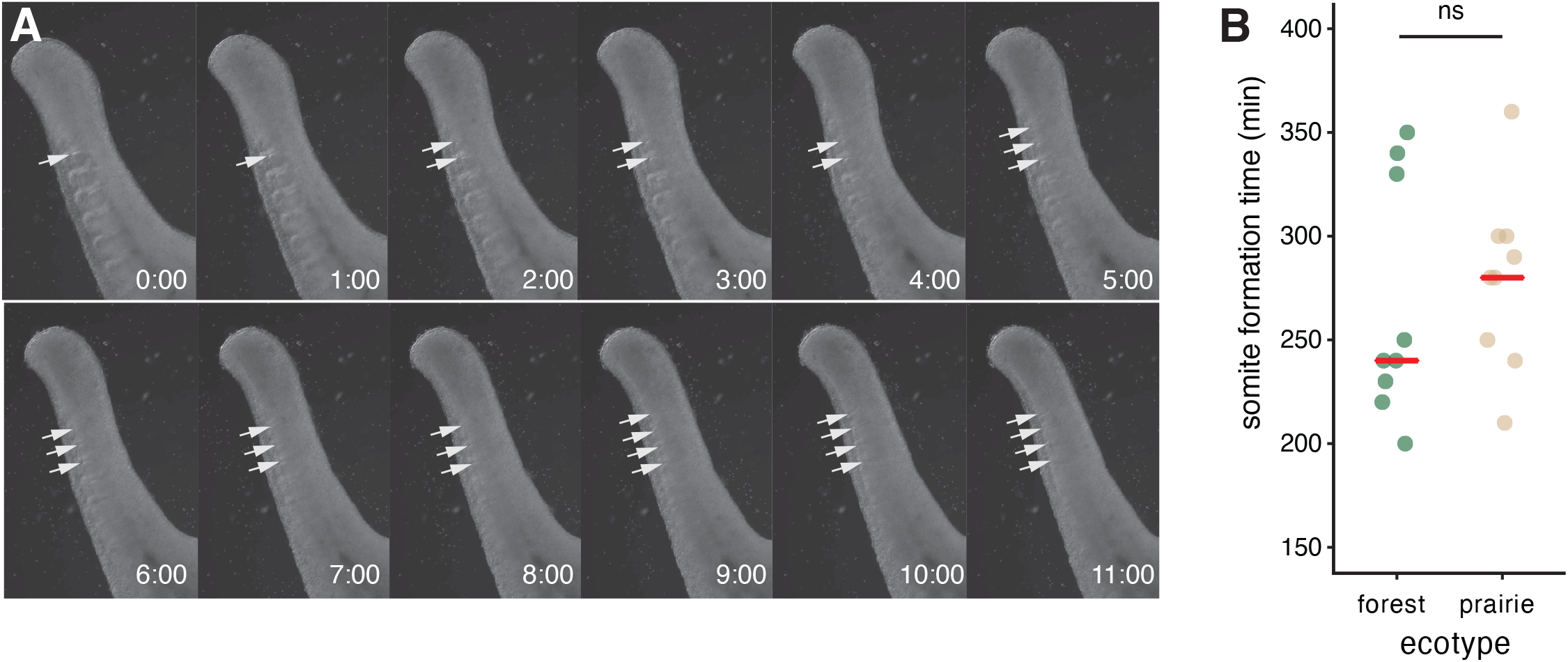
Timing of somite formation in embryonic tail explants. (Related to Fig. 6) **A**. Still images from time-lapse imaging of an E13.5 prairie embryonic tail explant. Recently formed somite borders are indicated by white arrows. Time stamps shown. **B**. Mean time between appearance of somite borders from tail explant cultures (forest, n = 9; prairie, n = 9; E12.5–15.5) shows no detectable difference in rate of somite formation (p = 0.45, Wilcoxon test).

**Figure S9.**
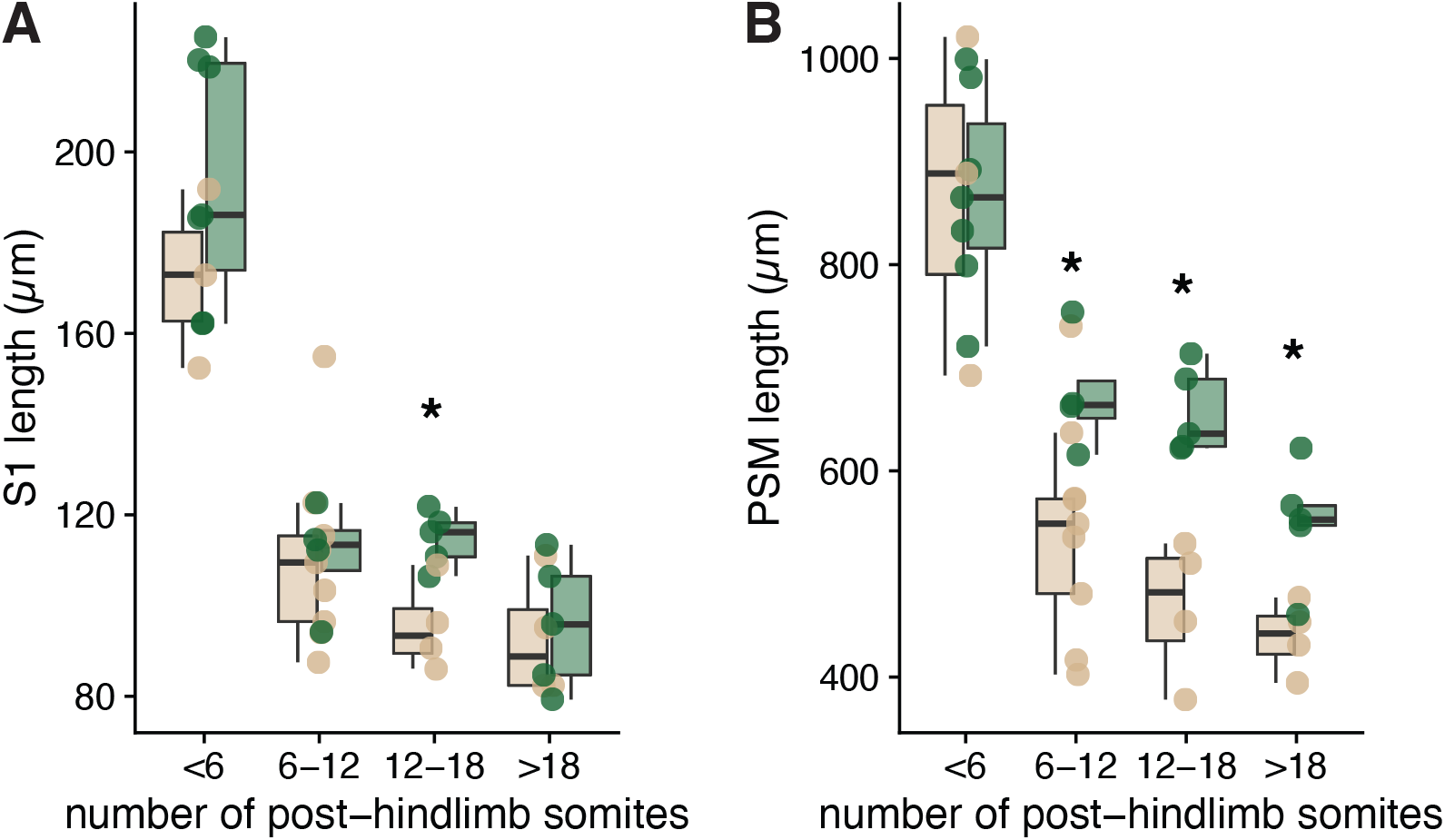
Binned somite- and PSM-length measurements. (Related to Fig. 6) **A**. Length of the most recently formed somite (S1) across embryonic tail development (E11.5–E15.5; as determined by number of post-hindlimb somites) measured in fixed specimens of forest (n = 20, green) and prairie (n = 18, tan) embryos. Data are the same as in Fig. 6A but in 6-somite bins. **B**. Length of presomitic mesoderm (PSM) measured in fixed specimens of forest (n = 20, green) and prairie (n = 18, tan) embryos. Data are the same as in Fig. 6A but in 6-somite bins. * = p < 0.05 (Wilcoxon test).

**Table S1.** Tail length QTL.

| trait | linkage group | 95% bayes conf interval | peak marker | position (cM) | LOD | p-value in multiple QTL model | PVE | mean additive effect (SE) | mean dominance effect (SE) |
| --- | --- | --- | --- | --- | --- | --- | --- | --- | --- |
| *tail total* | 3 | 47.2–80.6 | 562903406.374669r | 71.4 | 4.79 | 5e-3 | 1.7 | 0.96 (0.35) | 0.51 (0.48) |
| *tail total* | 8 | 22.3–38.9 | 562903868.1622618r | 23.5 | 5.6 | 0.02 | 1.2 | 1.02 (0.35) | 0.71 (0.49) |
| *tail total* | 14 | 0–16.6 | 562903608.62510r | 2.3 | 6.32 | 3e-7 | 5.1 | 1.82 (0.34) | 0.37 (0.50) |
| *tail total* | 17 | 58.3–64.6 | 562903716.995791r | 64.6 | 6.25 | 4e-5 | 3.3 | 1.16 (0.35) | 1.55 (0.49) |
| *tail total* | 20 | 76.2–84.7 | 562903388.1057339r | 84.2 | 8.27 | 1e-4 | 3.1 | 1.55 (0.37) | 0.02 (0.50) |
| *tail total* | X | 29.8–40.3 | 562903887.2705603r | 36.0 | 5.26 | 5e-5 | 4.5 | 0.4 (0.37) | N/A |
| *longest vert* | 13 | 42.7–64.2 | 562904146.173663r | 58.7 | 3.98 | 0.05 | 1.1 | 0.02 (0.02) | 0.03 (0.03) |
| *longest vert* | 17 | 56.7–68.6 | 562903898.92344r | 58.3 | 4.91 | 1e-3 | 1.9 | 0.04 (0.02) | 0.05 (0.03) |
| *longest vert* | 20 | 80.7–84.7 | 562903278.945341r | 84.7 | 13.52 | 2e-7 | 5.3 | 0.10 (0.02) | 0.01 (0.03) |
| *longest vert* | 21 | 35.8–67.8 | 562903066.344078r | 55.9 | 4.15 | 9e-3 | 1.7 | 0.05 (0.02) | -0.04 (0.03) |
| *longest vert* | X | 21.8–55.5 | 562903887.2705603r | 36.0 | 4.45 | 2e-4 | 4.0 | 0.02 (0.02) | N/A |
| *vert number* | 3 | 29.2–80.6 | 562903406.374669r | 71.4 | 3.76 | 2e-4 | 3.1 | 0.26 (0.06) | -0.06 (0.09) |
| *vert number* | 8 | 19.6–50.2 | 562903655.755532r | 29.5 | 5.1 | 6e-5 | 3.6 | 0.29 (0.06) | 0.05 (0.09) |
| *vert number* | 14 | 0–10.3 | 562903608.62510r | 2.3 | 5.06 | 3e-5 | 3.9 | 0.28 (0.06) | 0.11 (0.09) |

**Table S2.**
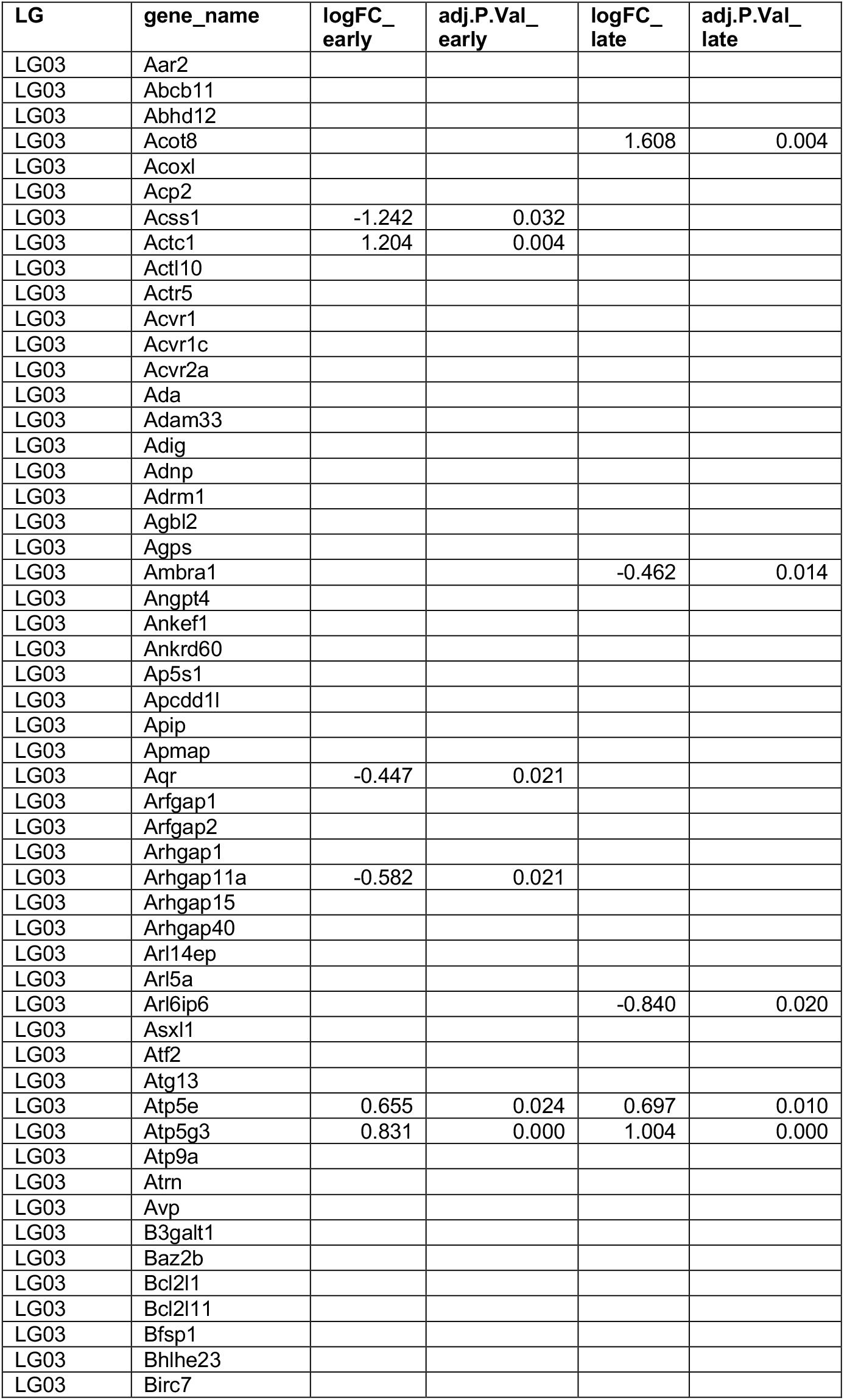
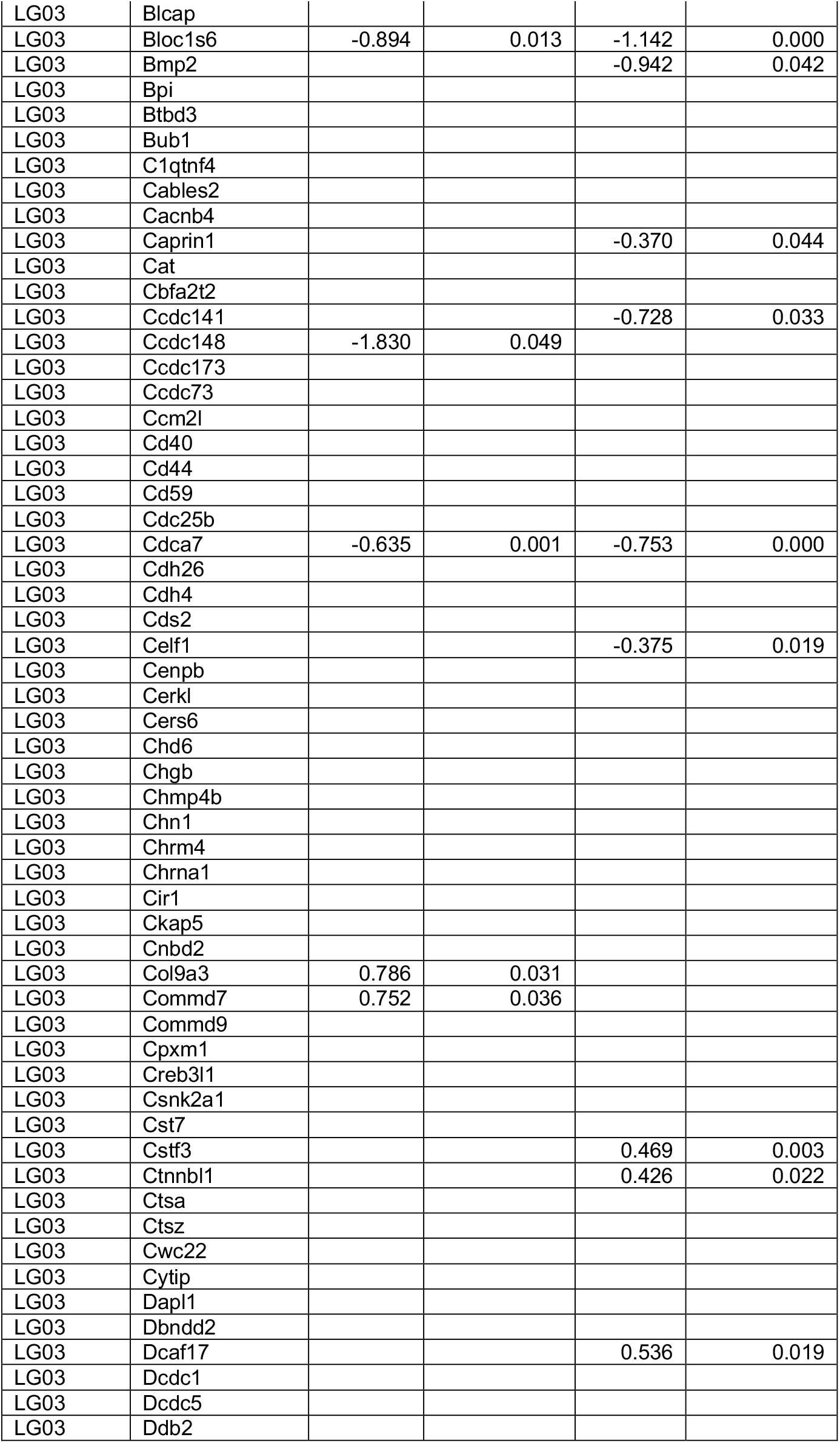
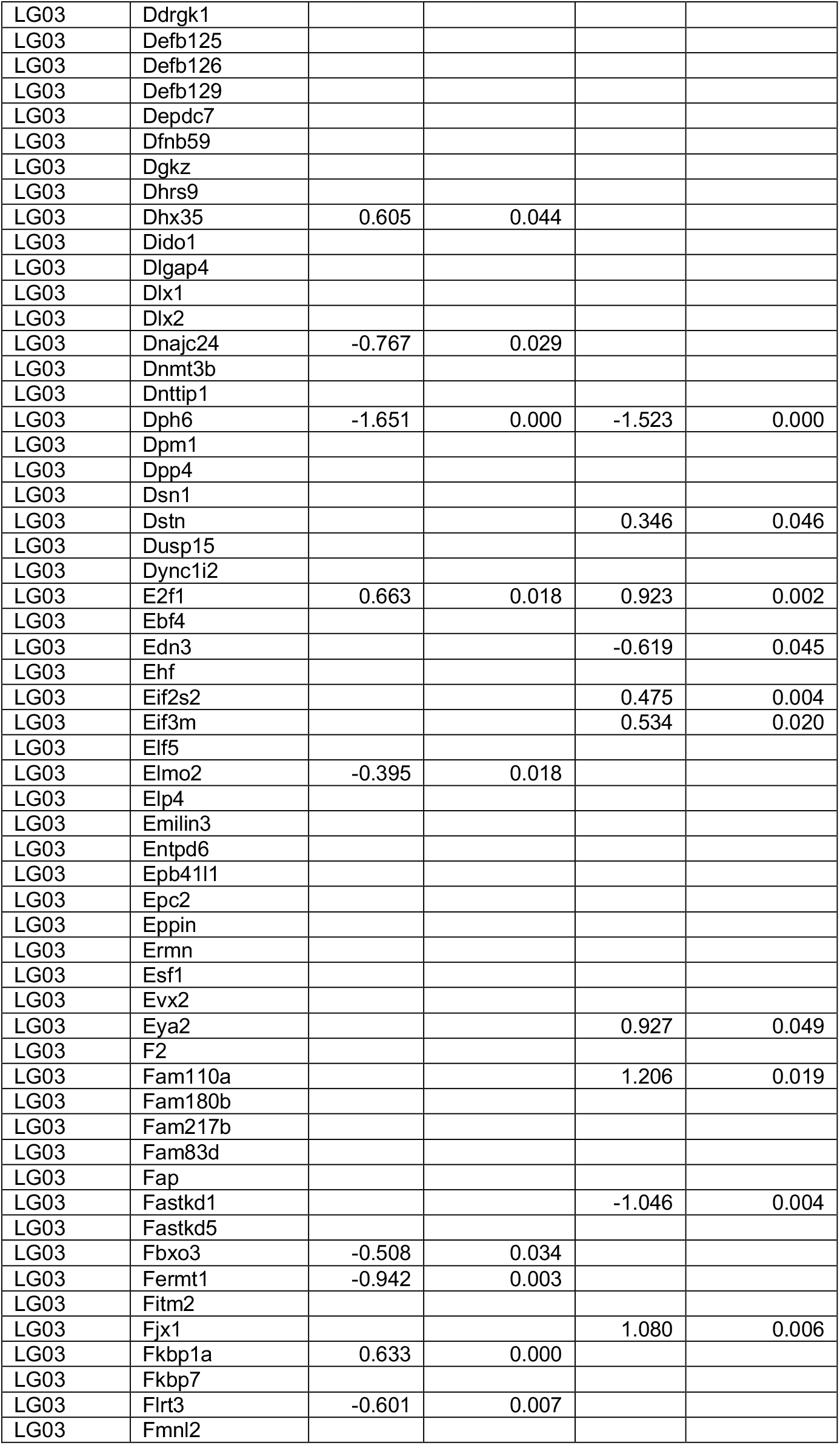
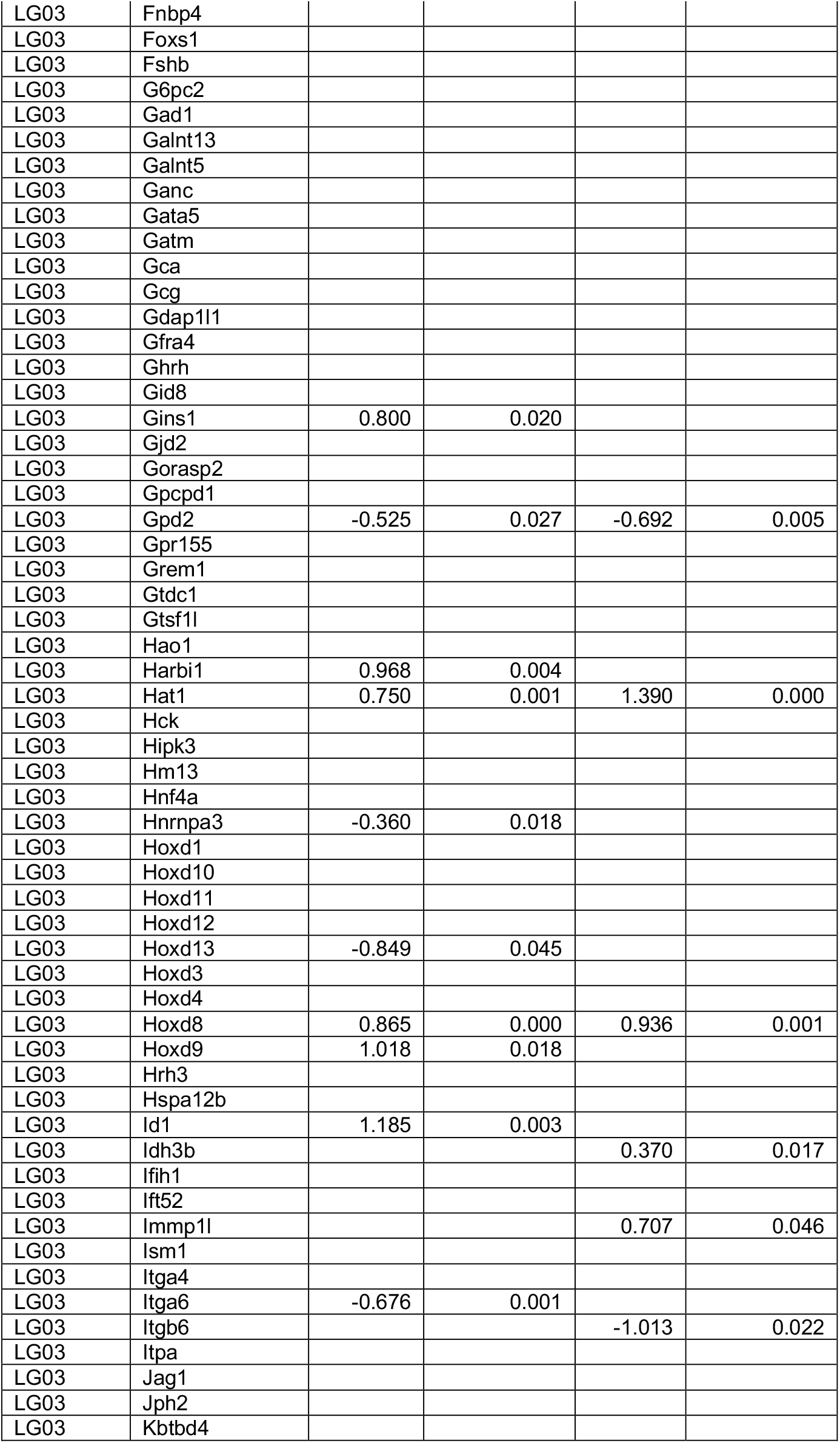
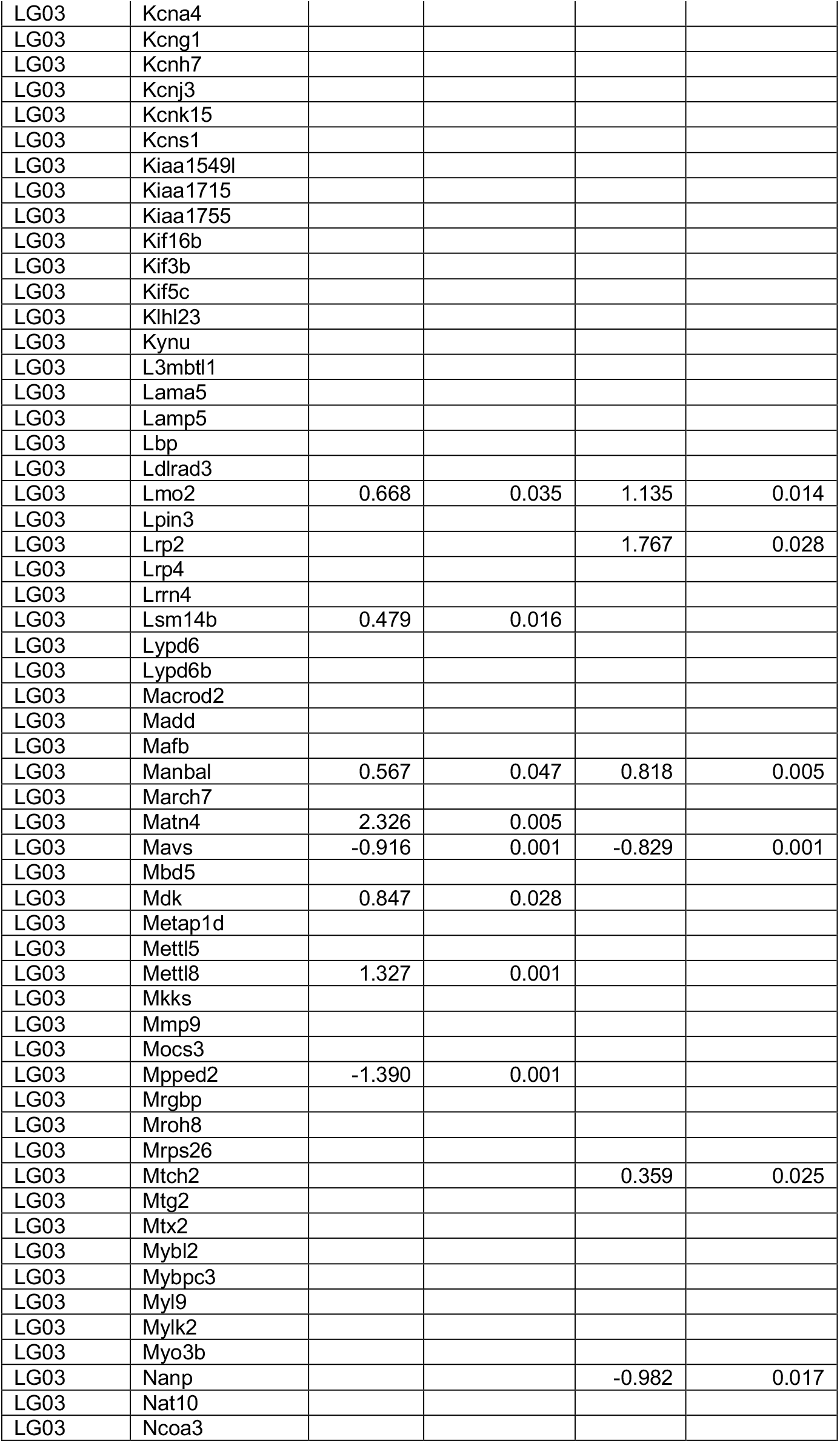
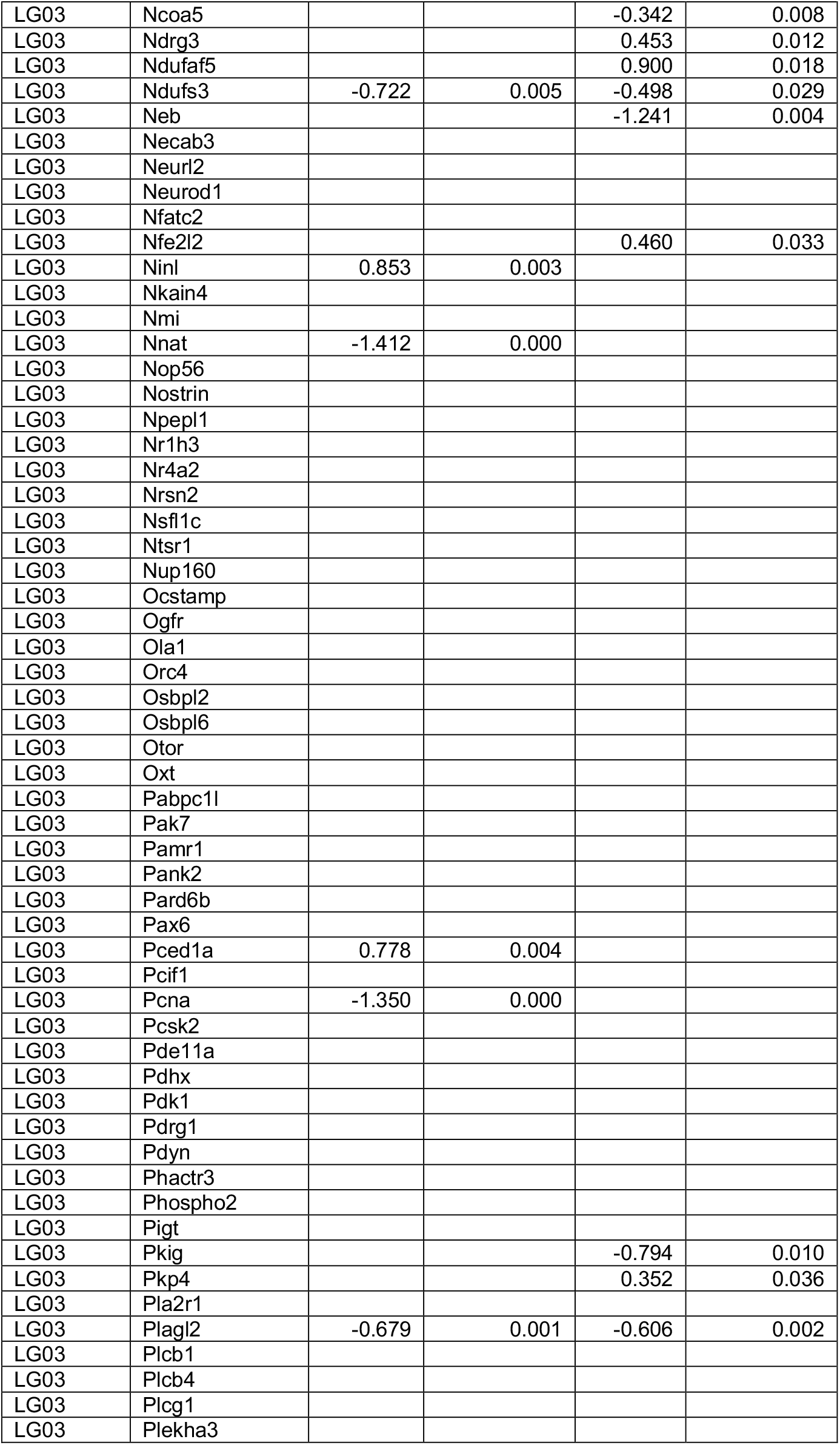
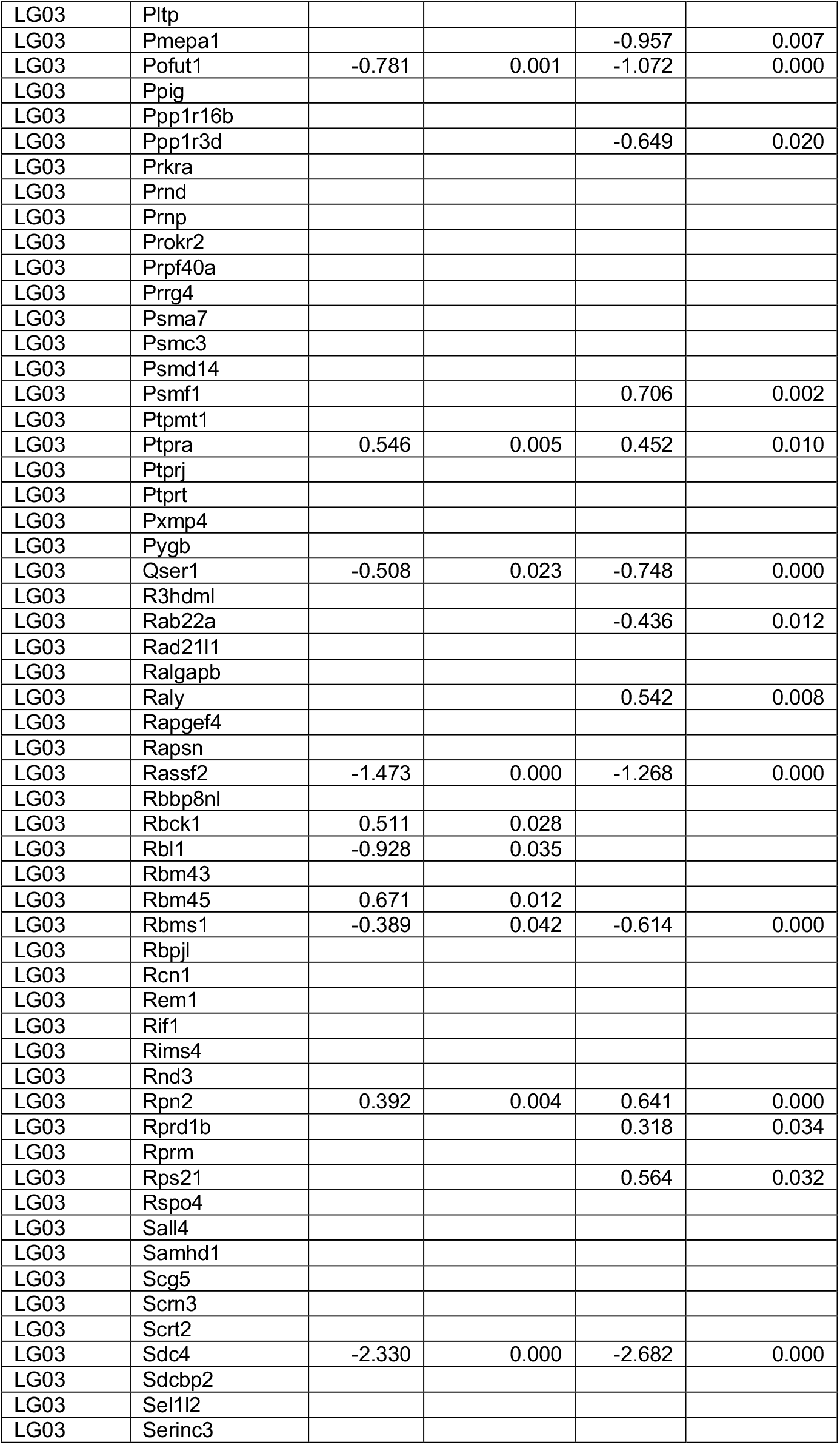
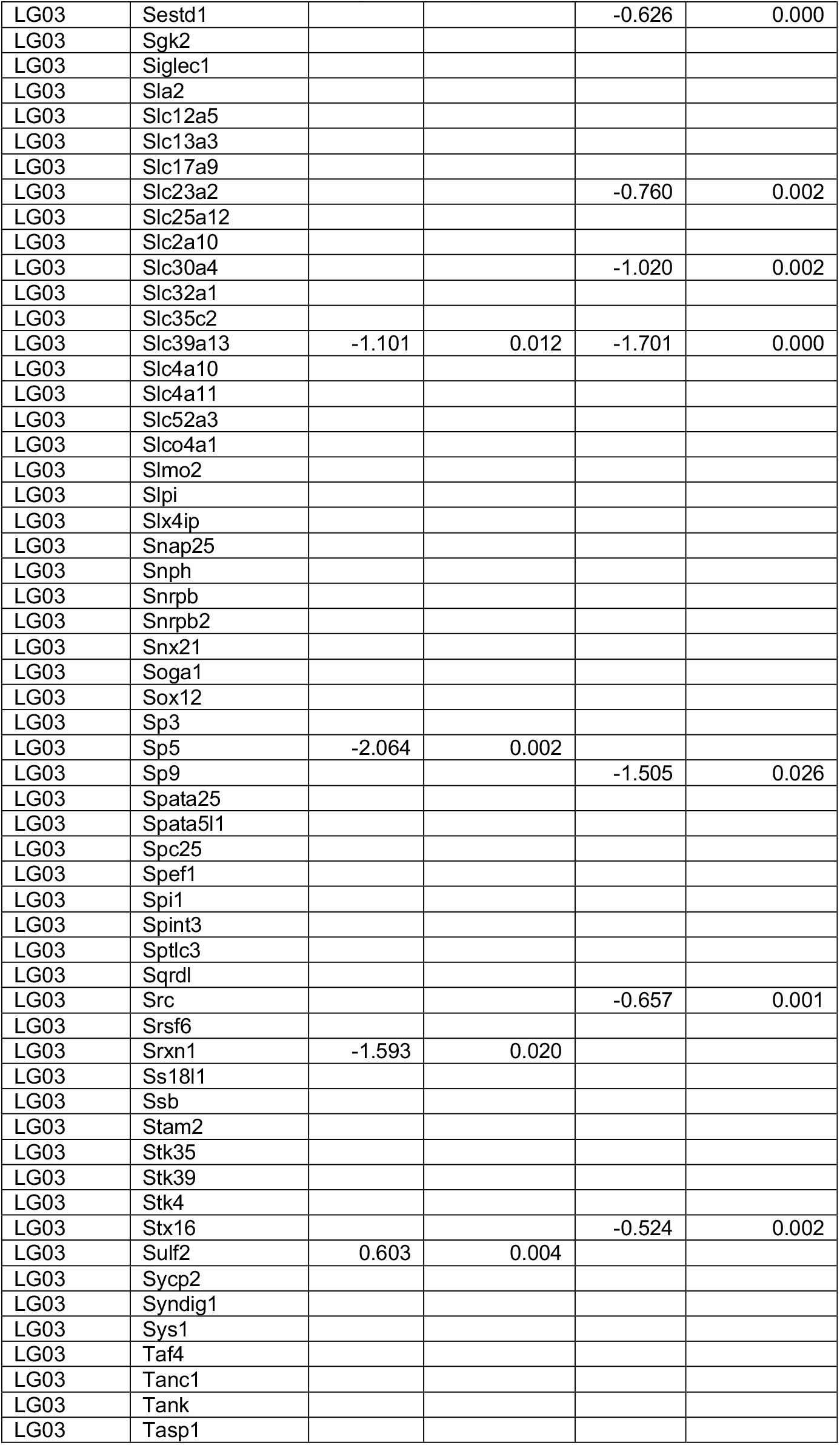
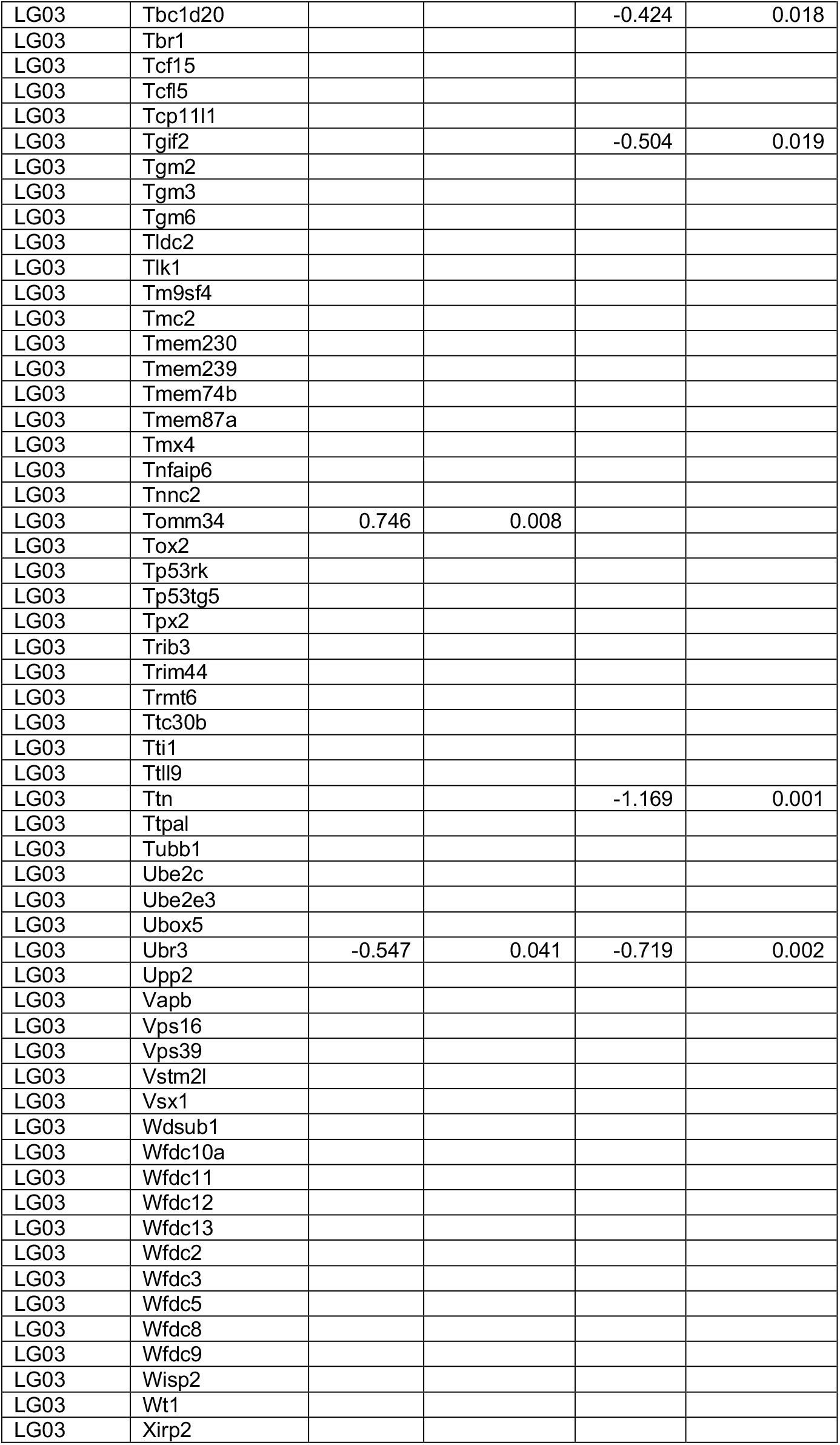
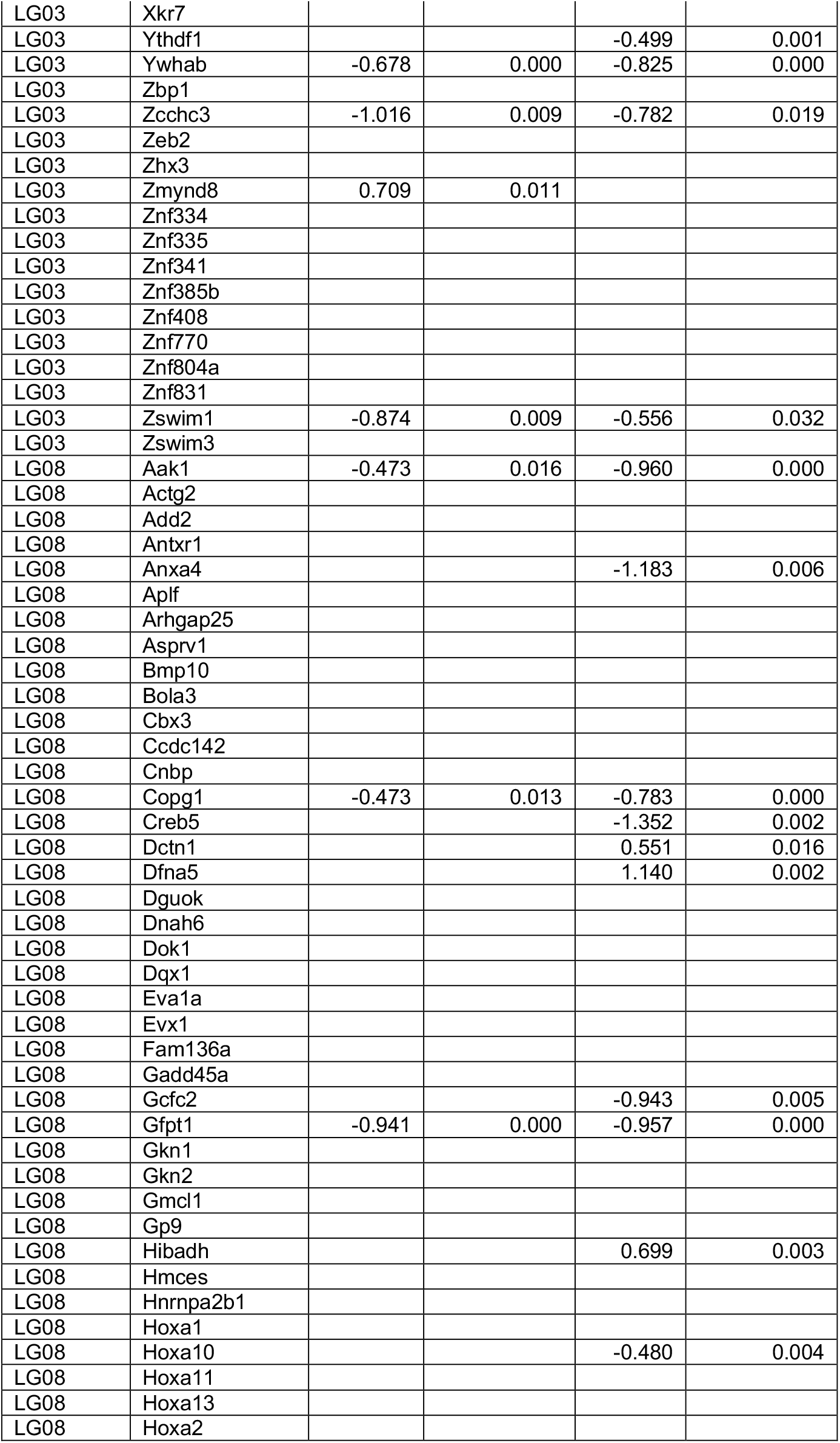
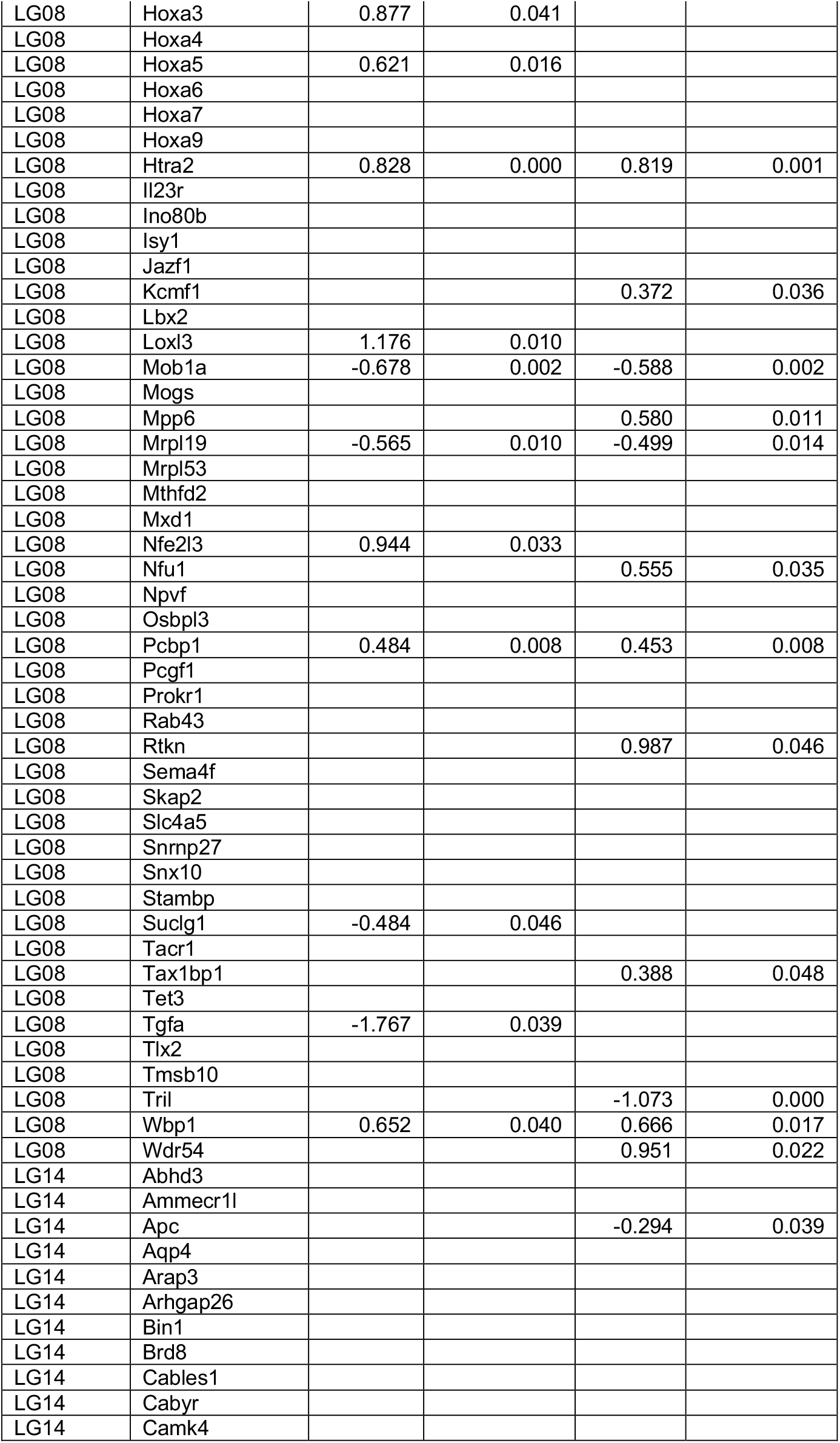
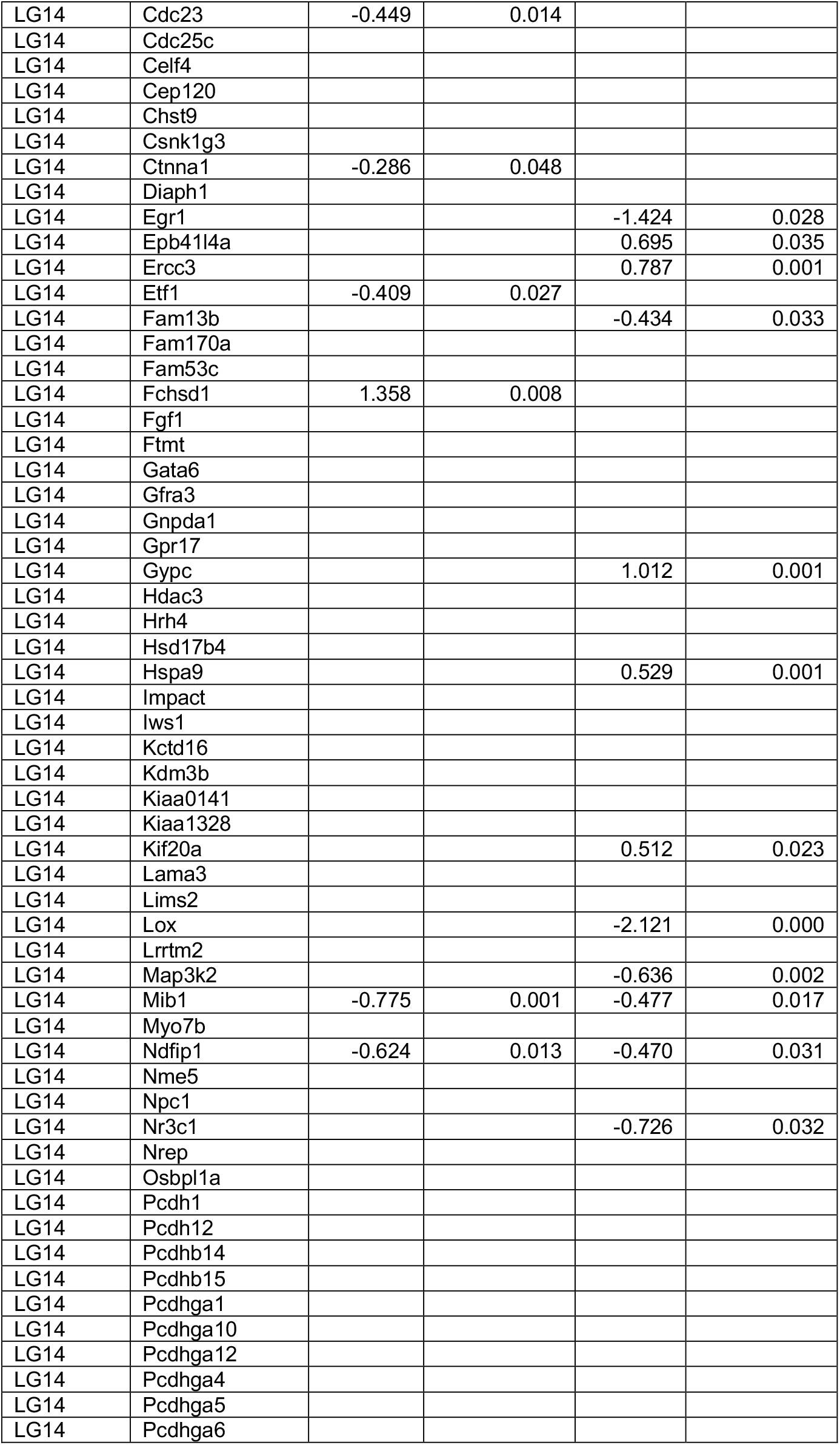
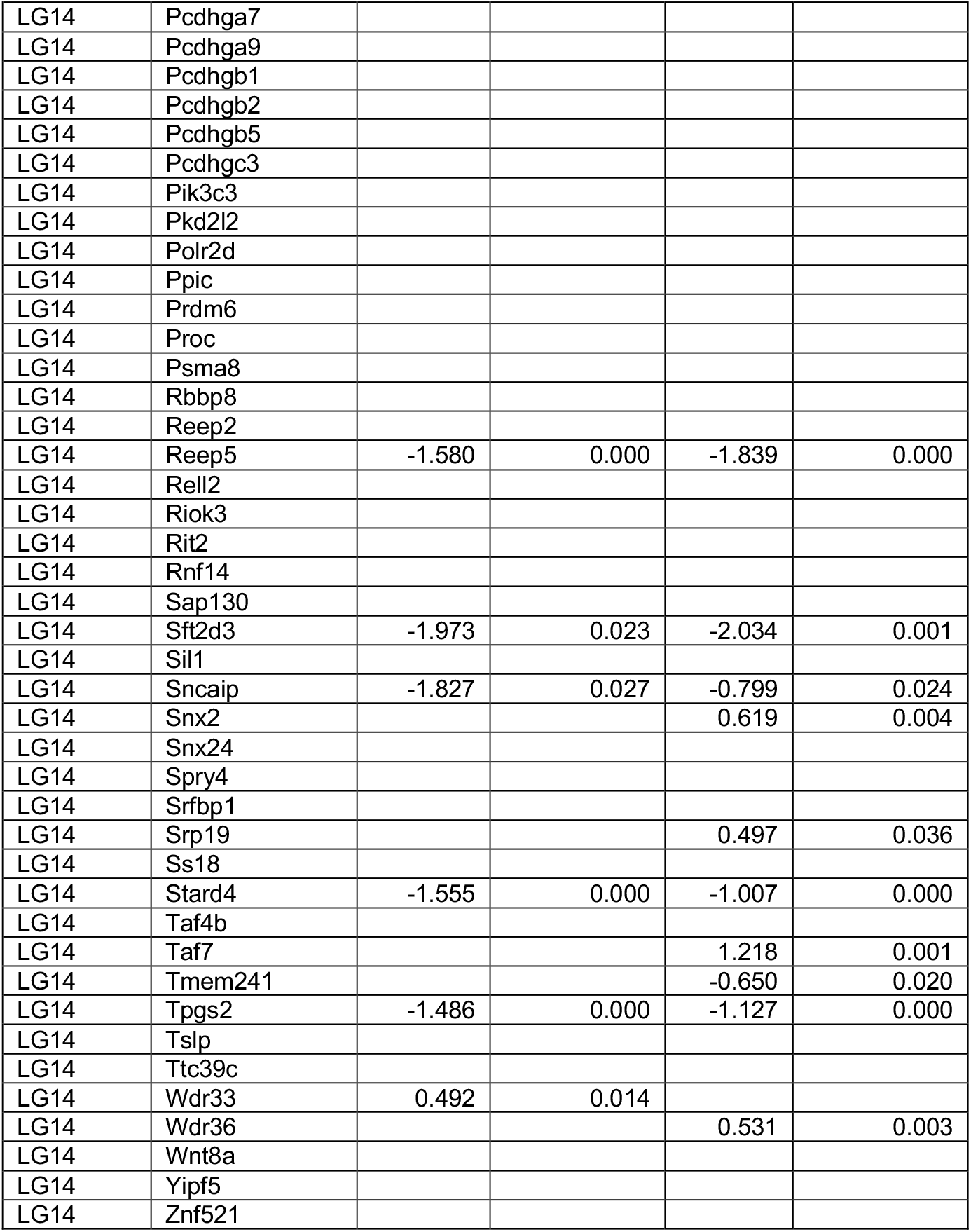
Annotated genes in QTL intervals with differential expression in embryonic tails.

**Table S3.** Phenotypes from Mouse Genome Informatics database.

| **MGI phenotype** |
| --- |
| abnormal_limb_mesenchyme_morphology |
| abnormal_vertebral_body_development |
| absent_apical_ectodermal_ridge |
| absent_caudal_vertebrae |
| absent_tail |
| decreased_caudal_vertebrae_number |
| decreased_ventral_ectodermal_ridge_size |
| elongated_metatarsal_bones |
| enlarged_tail_bud |
| increased_caudal_vertebrae_number |
| increased_tail_bud_apoptosis |
| long_limbs |
| long_tail |
| short_metatarsal_bones |
| short_tail |
| small_caudal_vertebrae |
| small_forelimb_buds |
| small_hindlimb_buds |
| small_tail_bud |
| thick_apical_ectodermal_ridge |
| thin_apical_ectodermal_ridge |
| truncated_tail_bud |
| vestigial_tail |

**Table S4.**
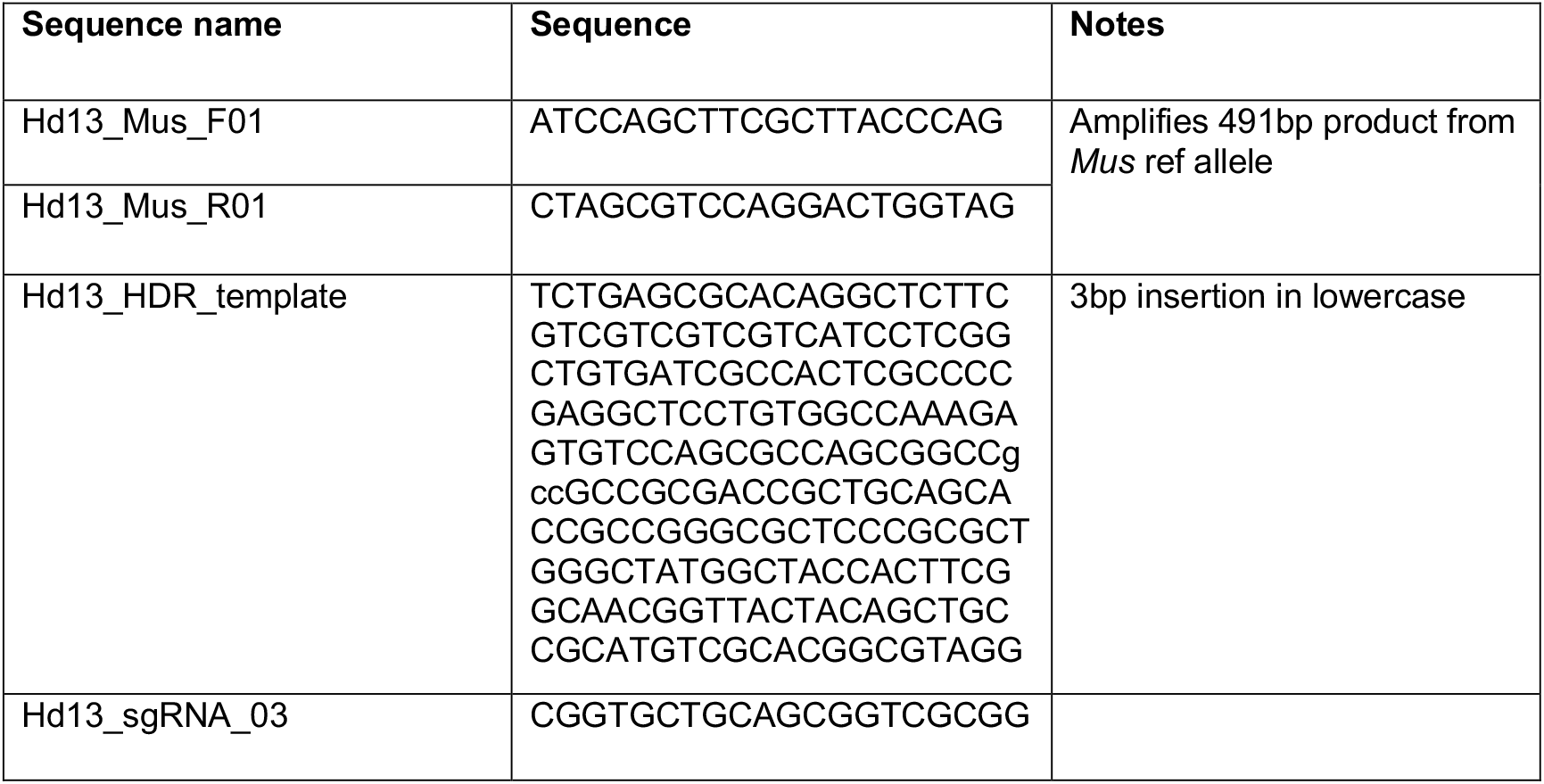
PCR primers, CRISPR HDR template and CRISPR guide sequences.

**Video S1**. Representative first cross attempt by a forest mouse (*P. maniculatus nubiterrae*)

**Video S2**. Representative first cross attempt by a prairie mouse (*P. maniculatus bairdii*)

